# Shifts in the human gut microbiome during cancer chemotherapy are diet-dependent

**DOI:** 10.64898/2025.12.23.696294

**Authors:** Kai R Trepka, Erin L Van Blarigan, Edwin F Ortega, Taylor Halsey, Summer D Bushman, Than S Kyaw, Christine A Olson, Gina Partipilo, Cecilia Noecker, Vaibhav Upadhyay, Daniel Gonzalez-Martinez, Chen Zhang, Daryll Gempis, Paige Steiding, Dalila Stanfield, Jessie A Turnbaugh, Alan P Venook, Eric P Skaar, Chloe E Atreya, Wesley A Kidder, Peter J Turnbaugh

## Abstract

Numerous studies have implicated both dietary intake and the human gut microbiome in colorectal cancer (CRC) treatment outcomes. However, little is known about how patients adjust their dietary intake during cancer chemotherapy or if these dietary changes contribute to treatment-associated alterations in the gut microbiome. We performed paired longitudinal diet and microbiome analysis during CRC treatment with oral fluoropyrimidines (NCT04054908) and validated key associations using cell culture assays. At each timepoint, diet was measured by averaging up to 3 consecutive days of 24-hour dietary records (NCI ASA24; 35 patients; 2.5±1.1 timepoints per patient), while microbiome composition was measured from stool (16S rRNA gene sequencing, metagenomics, qPCR; 40 patients; 5.6±1.4 timepoints per patient). Diet quality significantly decreased during chemotherapy. Carbohydrate and refined grain intake increased, accompanied by decreased consumption of fats, nuts and seeds, and fat-soluble micronutrients. Multiple individual dietary components were strongly linked to the gut microbiome. Decreases in theobromine intake correlated with decreases in overall microbial diversity and more gastrointestinal toxicities. Diet shifts partly explained changes in bacterial abundance during chemotherapy, including more severe depletion of *Faecalibacterium prausnitzii* in patients with decreased vitamin K_1_ intake. Changes in diet were correlated with multiple bacterial gene families involved in micronutrient metabolism and drug sensitivity. Increased copper intake was linked to decreased *Fusobacterium nucleatum* in patients and inhibited *F. nucleatum* in cell assays. Together, these data suggest that changes in diet during chemotherapy contribute to changes in gut bacterial diversity, taxonomic composition, and gene abundance. Our approach may generalize to other cancer therapies and emphasizes the need for collecting more robust dietary data in clinical microbiome studies.

## INTRODUCTION

The human gut microbiome is primarily shaped by environmental factors like food and drugs^1,2^. Host dietary intake changes the bacterial fitness landscape within the gastrointestinal tract, rapidly altering the abundance of gut microbial taxa and genes within days^3,4^. Similarly, pharmaceuticals, even those not traditionally considered antibiotics, can rapidly alter the gut microbiome^5–9^. In some cases, these drug-induced perturbations can persist for months to years after treatment cessation^10–12^. However, it remains unclear if and how dietary intake and medications interact to shape the human gut microbiome, and whether these interactions could have downstream consequences for treatment efficacy or side effect profiles.

Colorectal cancer (CRC) treatment provides a clinically relevant context to address these questions. Cancer chemotherapy often leads to gastrointestinal side effects that may lead individuals to dramatically shift their dietary intake^13^. However, few studies have comprehensively and longitudinally quantified dietary intake during CRC treatment^14^, with existing studies focusing on malnourishment and macronutrients^15^. These gaps in knowledge have limited the ability to make informed dietary recommendations or to propose specific dietary interventions for patients during treatment aimed at improving treatment outcomes^16^.

There is also an extensive and rapidly growing literature on the impact of cancer chemotherapy on the gut microbiome and, in turn, the ability of the microbiome to influence treatment outcomes^7,17^. For example, we recently identified multiple gut bacterial chemoprotective genes that increase in abundance during oral fluoropyrimidine treatment, rescuing drug toxicity by contributing to drug clearance^18^ and producing vitamin K_2_^11^. Studies in cell cultures suggested that fluoropyrimidines directly impact gut microbial community structure due to their ability to inhibit the growth of select taxa^18,19^; however, whether concomitant changes in host dietary intake to contribute to the observed changes in the gut microbiome remains an open question.

Herein, we describe an integrated analysis building on our published metagenomic sequencing data from 40 CRC patients over the course of oral fluoropyrimidine chemotherapy (NCT04054908)^18^ together with longitudinal data on host dietary intake. We show that host dietary intake changed during cancer treatment and explained shifts in gut microbiota diversity, composition, and functional potential. These observational data were then used to design controlled experiments in cell culture that support a causal role of individual nutrients in altering the abundance of gut bacterial species and genes relevant to treatment outcomes. Taken together, our results (*i*) reinforce the importance of host diet in shaping drug-microbiome interactions; (*ii*) emphasize the need to account for diet in clinical microbiome studies; and (*iii*) provide a foundation to design dietary interventions to optimize cancer treatment outcomes.

## RESULTS

We conducted the gut microbiome and oral fluoropyrimidine (GO) clinical study (ClinicalTrials.gov NCT04054908), a prospective longitudinal study investigating the impact of oral fluoropyrimidines on diet and the gut microbiome^18^. Stool was collected at up to 7 time points spanning pretreatment to after cycle 3 of chemotherapy (**Fig. 1a**). Three-day dietary records were collected over the 3 days prior to cycle 1 (C1), cycle 2 (C2), and cycle 3 (C3), with dietary quantities adjusted for total caloric intake prior to analysis (*Methods*). Of the 52 patients enrolled, 40 submitted at least 1 stool sample, of which 35 submitted at least 1 valid set of dietary records (**Extended Data Fig. 1**). Patients with dietary information were distributed among 3 subcohorts (**Fig. 1a, Table S1**): (subcohort A) CAP as monotherapy or part of a standard-of-care regimen (*n*=20), (subcohort B) TAS-102 (trifluridine/tipiracil, *n*=6), and (subcohort C) CAP with bevacizumab and pembrolizumab immunotherapy (patients participating in ClinicalTrials.gov ID NCT03396926, *n*=9). These 35 participants had a mean age of 53.0±11.1 years; 51% were male, 83% identified as white, and 6% identified as Hispanic or Latino (**Table S1**). 83% of patients were in tumor, node, metastasis (TNM) stages III (51%) and IV (31%), and 66% had prior surgery (**Table S1**). We pooled patients across the 3 treatment subcohorts to increase statistical power, with post hoc examination of key findings stratified by cohort.

**Figure 1:**
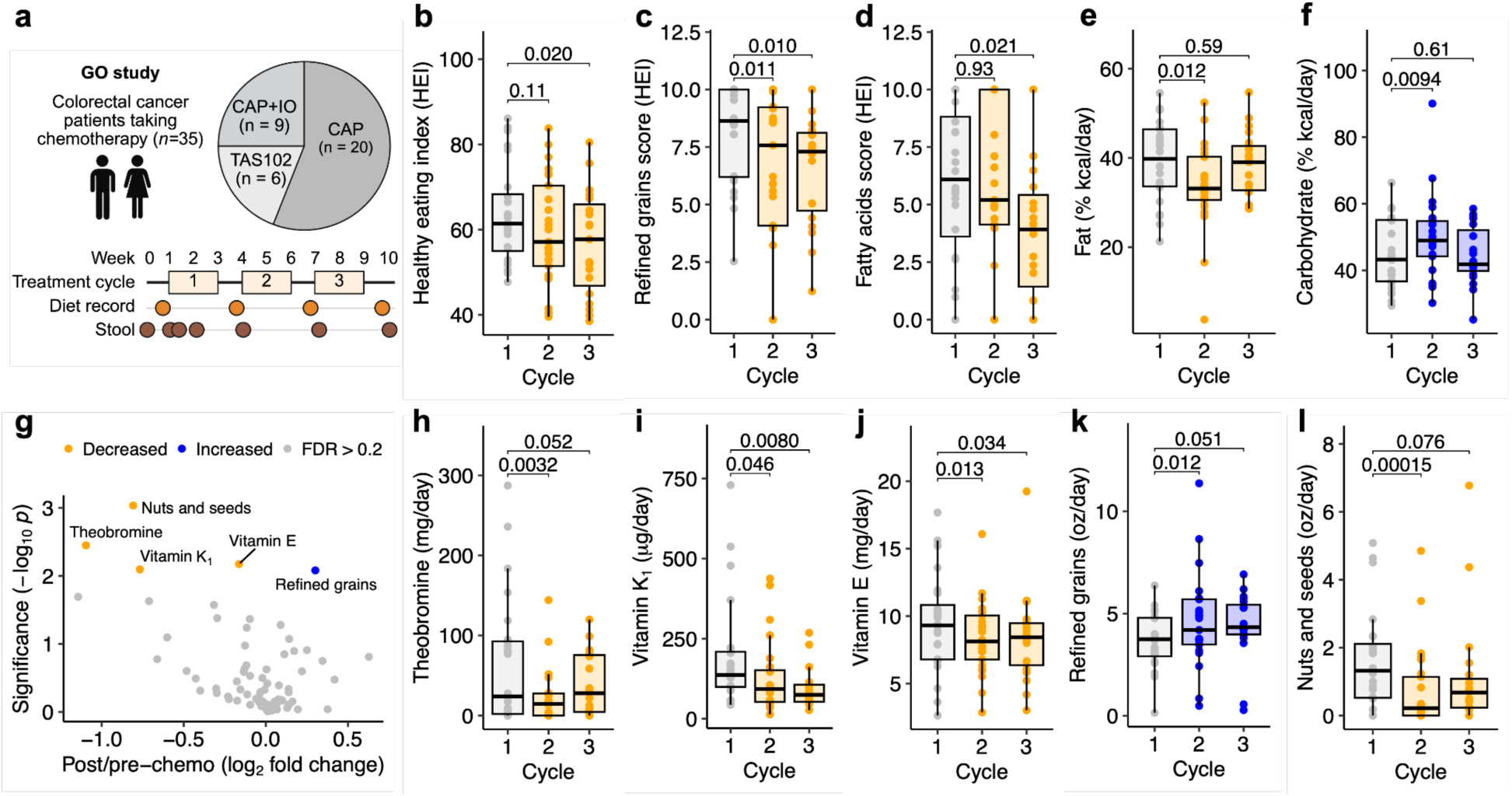
Dietary intake changes in patients with colorectal cancer during oral chemotherapy. (**a**) Gut microbiome and Oral fluoropyrimidine (GO) study design depicting patients treated with capecitabine (CAP), TAS102, or combination CAP + immunotherapy (IO), with dietary surveys conducted in the 3 days leading up to each treatment cycle. (**b-f**) Change in healthy eating index (HEI) (**b**), HEI refined grains and fatty acids scores (**c,d**), and macronutrient intake (**e,f**) during the study. *p-*values: Wilcoxon signed-rank test. (**g**) Volcano plot of ASA24 diet survey variables with respect to treatment time. Points represent increased (blue), decreased (orange), or non-significant (grey) dietary variables after treatment (Benjamini-Hochberg false discovery rate [FDR] < 0.2). (**h-l**) Change in intake of significant dietary variables from (**g**) during the study. *p*-values: Wilcoxon signed-rank test (**b-f,h-l**), mixed-effects model, Intake ∼ PrePost + 1|Patient (**g**). *n*=68 samples from 25 patients.

We assessed diet quality during treatment using the healthy eating index (HEI-2020)^20^. HEI was significantly decreased during chemotherapy (**Fig. 1b**). We investigated each of the 13 HEI components, finding significantly decreased refined grains score (*i.e.* increased refined grains consumption, **Fig. 1c**) and fatty acids score (*i.e.* decreased unsaturated:saturated fatty acids ratio, **Fig. 1d**). The decreased refined grains and fatty acids scores were sufficient to explain the decreased HEI following chemotherapy (**Extended Data Fig. 2a**). We observed similar decreases in the Alternative Healthy Eating Index^21^ (**Extended Data Fig. 2b-d**). Macronutrient intake also changed significantly at C2, with decreased fat (**Fig. 1e**) and increased carbohydrate intake (**Fig. 1f**). Protein and total energy intake did not change throughout treatment (**Extended Data Fig. 2e,f**).

Next, we sought to identify which individual dietary features varied during treatment. After dereplication of highly correlated diet variables (**Fig. S1a, Tables S2, S3**) and false discovery correction of 79 tested dietary features (**Table S3**), we identified 3 micronutrients and 2 food groups that changed following chemotherapy (**Fig. 1g**). Patients consumed fewer fat-soluble micronutrients, with significant decreases in theobromine (**Fig. 1h**), vitamin K_1_ (**Fig. 1i**), and vitamin E (**Fig. 1j**). The proportion of patients meeting guideline-recommended intake levels for vitamin K_1_ and vitamin E declined markedly (K_1_ [C1: 72%; C2: 44%]; E [C1: 16%; C2: 0%]; **Table S4**). Refined grain intake was significantly increased (**Fig. 1k**). Nut and seed intake was significantly decreased (**Fig. 1l**), which was sufficient to explain decreased fat and vitamin E intake at C2 (**Extended Data Fig. 2g-j**). These dietary shifts were similar across all 3 subcohorts (CAP, TAS102, CAP+IO; **Extended Data Fig. 2k**) and across patient metadata categories (BMI, sex, age, stage, tumor sidedness; **Extended Data Fig. 2l**), despite considerable patient-to-patient variation (**Fig. S1b,c**).

Baseline diet was associated with treatment outcomes. Baseline total meat, poultry, and seafood intake was significantly lower in patients who experienced progressive disease (**Extended Data Fig. 3a,b**). Further, we found significantly lower baseline carbohydrate, total sugar, added sugar, and starchy vegetable intake in patients who required dose delays or reductions (**Extended Data Fig. 3c-g**). Finally, baseline theobromine intake was significantly lower in patients who experienced hand-foot syndrome (HFS) in our cohort (**Extended Data Fig. 3h,i**) and in an independent cohort of CAP-treated CRC patients^11^ (**Extended Data Fig. 3j**). These results were robust to tested patient metadata variables (sex, BMI category, tumor sidedness, CRC onset before age 50 [young-onset CRC, yoCRC], stage); all tested metadata variables were not significantly associated with progressive disease, dose adjustments, or HFS (Fisher’s exact *p* > 0.05 for all). No metadata variables were significantly associated with baseline intake of meat/poultry/seafood, carbohydrate, total sugar, added sugar, starchy vegetable, or theobromine (Welch’s *t*-test FDR > 0.2 for all). While adjustment for sex slightly attenuated the relationship between starchy vegetables and dose delays or reductions, all other observed diet-outcome relationships were not weakened by metadata adjustment (**Table S5**).

Next, we evaluated diet-taxon interactions during chemotherapy with 16S rRNA gene sequencing. We combined our diet surveys with a subset of our published GO 16S dataset^18^, resulting in an integrated diet-16S dataset of 85 samples from 35 patients with 71,502 ± 7,872 high-quality reads per sample (**Table S6**). To leverage the longitudinal nature of this combined dataset, we correlated the change in dietary intake with change in microbial diversity metrics, computing only changes between consecutive within-patient timepoints to control for baseline interpatient variability in starting diet and microbiota (*Methods*). We performed this analysis for all 79 dereplicated diet variables (**Table S3**).

While multiple diet variables showed interesting trends, only theobromine was significant after multiple hypothesis correction (**Fig. 2a**). Increases in theobromine intake were positively correlated with increases in the number of observed genera (**Fig. 2b**) and the Shannon index of diversity (**Extended Data Fig. 4a,b**). Theobromine and diet quality were among 5 diet variables significantly associated with changes in microbiota composition (**Fig. 2c-h**). Interestingly, gastrointestinal toxicities were associated with both baseline microbiome composition (**Extended Data Fig. 4c**) and with greater decreases in theobromine intake (**Fig. 2i**).

**Figure 2:**
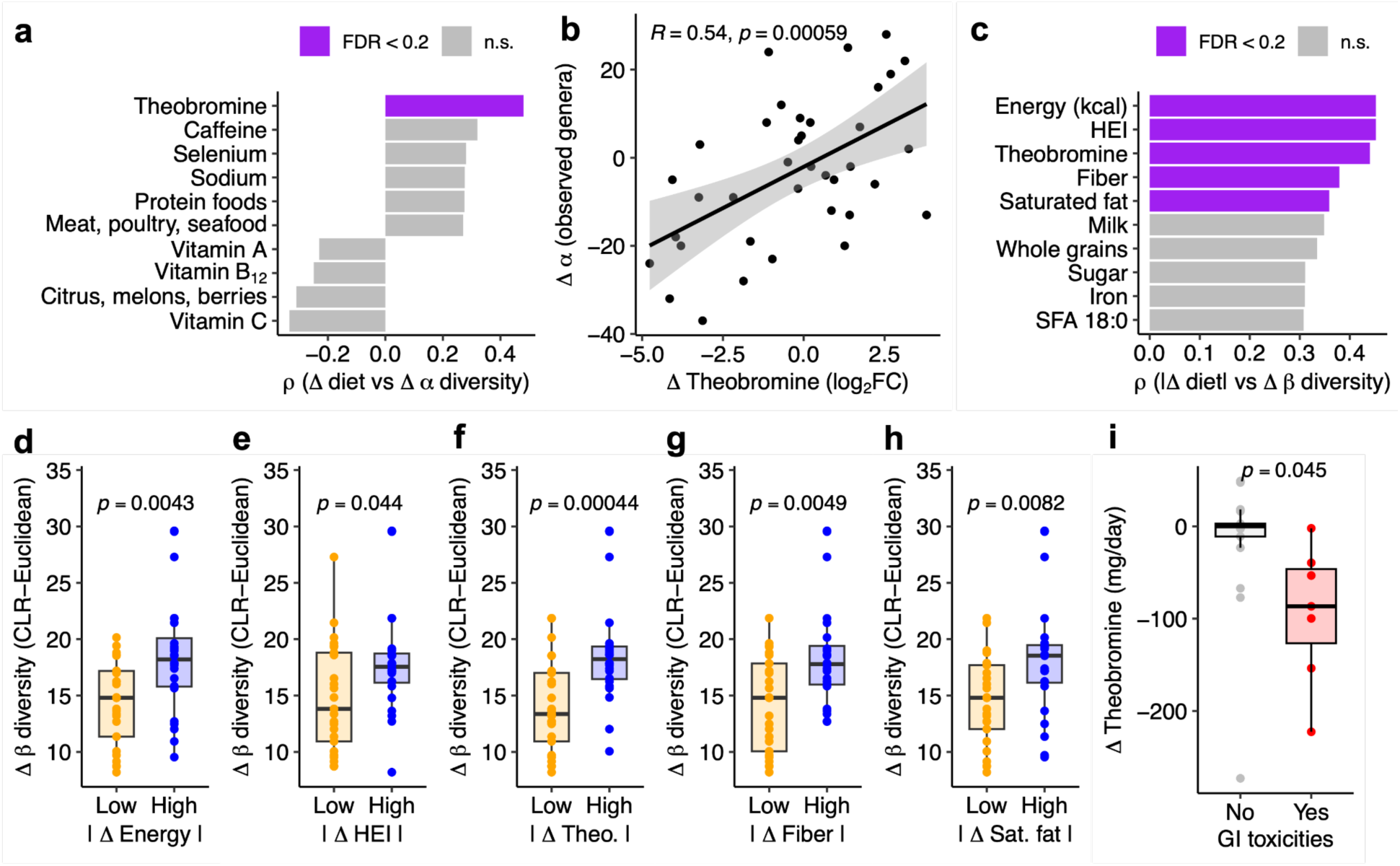
Dietary intake dynamics correlate with gut microbiota diversity shifts. (**a**) Top 10 diet variables associated with shifts in alpha diversity (number of observed genera). (**b**) Change in alpha diversity (number of observed genera) vs change in theobromine intake. (**c**) Top 10 diet variables associated with shifts in beta diversity (CLR-Euclidean transformed genera composition). (**d-h**) Change in beta diversity vs absolute value of change in dietary intake for significant dietary variables from (**c**), where Low/High indicate below/above median absolute intake shift. (**i**) Change in theobromine intake (C2 minus C1) vs presence of gastrointestinal (GI) toxicities. *p*-values: (**a,c**) Benjamini-Hochberg false discovery rate (FDR)-adjusted Spearman correlation for change in rank-transformed dietary intake vs change in rank-transformed diversity [signed for (**a**), absolute value for (**c**)] between each pair of consecutive within-patient timepoints, with FDR > 0.2 called as not significant (n.s.); (**b**) Linear regression of log₂ fold-change of dietary intake vs log₂ fold-change of diversity between each pair of consecutive within-patient timepoints, with solid black line indicating the line of best fit and grey bands the 95% confidence interval; (**d-i**) Mann-Whitney U test. *n*=48 Δdiet-Δdiversity pairs (**a,b**); 51 Δdiet-Δdiversity pairs (**c-h**); 24 subjects with paired diet C1/C2 (**i**).

We further investigated theobromine due to its associations with multiple aspects of the gut microbiota and side effect profiles (**Fig. 2, Extended Data Fig. 3h-j**). In our cohort, chocolate desserts and tea were the most common sources of theobromine (**Extended Data Fig. 4d**), with chocolate desserts accounting for the majority of theobromine consumption (**Extended Data Fig. 4e**). Chocolate dessert-derived theobromine was sufficient to explain theobromine-diversity relationships across subcohorts (**Extended Data Fig. 4f-l**) and the theobromine-GI toxicity relationship (**Extended Data Fig. 4m,n**).

Diet shifts partially explained the dynamics of chemotherapy-altered gut bacteria. We correlated change in diet with change in the abundance of 15 prevalent amplicon sequence variants (ASVs) that we previously identified as altered during chemotherapy^18^ (**Table S7**). This analysis identified 8 diet-ASV interactions (**Fig. 3a**), including 2 positive and 6 negative interactions encompassing 5 unique ASVs (**Fig. 3a, Extended Data Fig. 5a-f, Table S7**).

**Figure 3:**
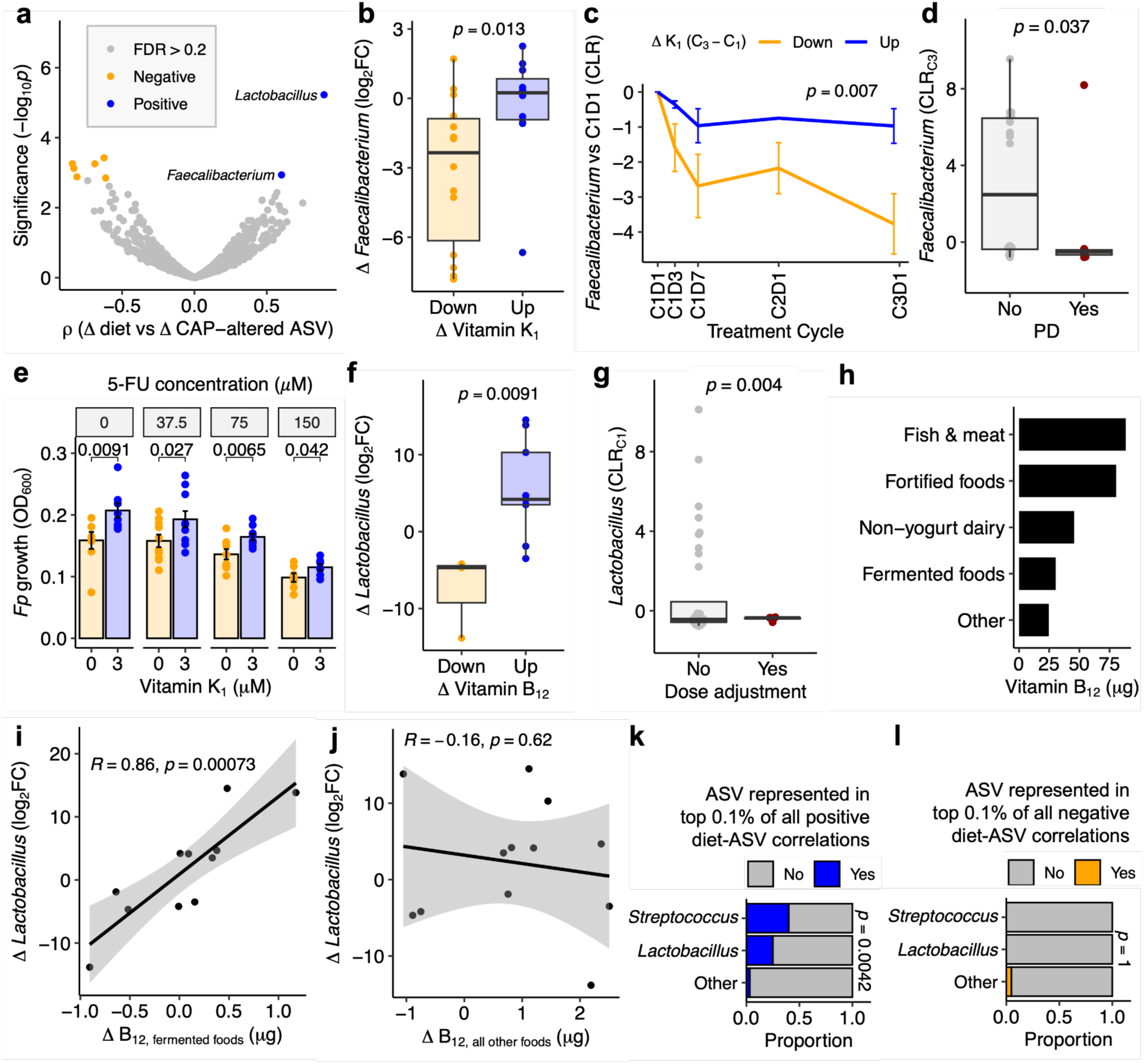
Altered dietary intake is associated with gut bacterial dynamics during chemotherapy. (**a**) Volcano plot of interactions between diet variables (food groups, macronutrients, and micronutrients) and 15 amplicon sequence variants (ASV) significantly altered during chemotherapy^18^. Each point represents a single diet-ASV pair. (**b**) Change in *Faecalibacterium* ASV abundance vs change in vitamin K_1_ intake. Each point represents the difference between 2 consecutive timepoints within a single patient. (**c**) Change in *Faecalibacterium* ASV abundance stratified by the change in vitamin K_1_ intake between C1 and C3. (**d**) *Faecalibacterium* ASV abundance during C3 vs progressive disease (PD) status. (**e**) Carrying capacity (OD_600_) of *Faecalibacterium prausnitzii* WB-STR-0032 (*Fp*) grown for 24 hours in BHI^CHV^ ± 5-FU (0, 37.5, 75, 150 μM) and vitamin K_1_ (0, 3 μM). Vitamin K_1_ promotes *Fp* growth in vitro (*n*=7-10 replicates pooled from 2 independent experiments). (**g**) Change in *Lactobacillus* ASV abundance vs change in vitamin B_12_ intake. Each point represents the difference between 2 consecutive timepoints within a single patient. (**f**) Maximum *Lactobacillus* ASV abundance during C1 vs dose adjustment status. (**h**) Barplot of vitamin B_12_ source. (**i,j**) Scatterplot of changes in fermented food-derived (**i**) and non-fermented food-derived (**j**) vitamin B_12_ vs changes in *Lactobacillus* ASV abundance. (**k,l**) Food-derived genera *Streptococcus* and *Lactobacillus* are overrepresented in the top 0.1% of positive Δdiet-ΔASV correlations (**k**), but not in the top 0.1% of negative Δdiet-ΔASV correlation correlations (**l**). *p*-values: (**a**) Spearman correlation of change in rank-transformed dietary intake vs change in rank-transformed central log ratio (CLR)-normalized ASV abundance, with Benjamini-Hochberg false discovery rate (FDR) correction across all tested diet-ASV pairs; (**b,g**) Mann-Whitney U test; (**c**) Group (diet) term from two-way ANOVA; (**d,g**) one-tailed Welch’s *t*-test; (**e**) one-tailed Welch’s *t*-test; (**i,j**) Linear regression of log₂ fold-change of dietary intake vs log₂ fold-change ASV abundance between each pair of consecutive within-patient timepoints, with solid black line indicating the line of best fit and grey bands the 95% confidence interval; (**k,l**) Fisher’s exact test.

Decreases in vitamin K_1_ intake were linked with decreases in *Faecalibacterium prausnitzii* (**Fig. 3b**), with similar trends observed across all 3 sub-cohorts (**Extended Data Fig. 5g-i**). *F. prausnitzii* was depleted during treatment, with significantly larger declines observed in patients who decreased their vitamin K_1_ intake (**Fig. 3c**). Low *F. prausnitzii* abundance at cycle 3 was associated with disease progression (**Fig. 3d**). We grew *F. prausnitzii* WB-STR-0032 in BHI^CHV^ ± 3 μM vitamin K_1_ ± active CAP metabolite 5-FU (37.5, 75, 150 μM). We found vitamin K_1_ promotes *F. prausnitzii* growth irrespective of 5-FU (**Fig. 3e**).

*Lactobacillus fermentum* was linked with increased vitamin B_12_ intake (**Fig. 3f**) and fewer dose adjustments (**Fig. 3g**). Between cycle 1 and cycle 2, *L. fermentum* was enriched exclusively in patients with increased vitamin B_12_ (**Extended Data Fig. 6a**). Because *L. fermentum* is commonly found in yogurt and fermented vegetables^22^, we wondered whether *L. fermentum* and other bacteria might be directly inoculated from food. Though fermented foods accounted for a minority of B_12_ intake (**Fig. 3h**), fermented food-derived B_12_ explained *L. fermentum* dynamics (**Fig. 3i,j**). We then correlated change in diet with change in ASV abundance for each of 258 prevalent ASVs (*n*=22,430 diet-ASV interactions). Out of the 258 tested ASVs, *Streptococcus* and *Lactobacillus* were overrepresented among the top 0.1% of positive diet-ASV interactions (**Fig. 3k**), including a significant yogurt-*Streptococcus thermophilus* relationship (**Extended Data Fig. 6b,c**). *Streptococcus* and *Lactobacillus* spp. were notably absent from the top 0.1% of negative diet-ASV interactions (**Fig. 3l**).

Next, we used metagenomic sequencing to assess diet-gene interactions during treatment. We combined our diet surveys with a subset of our published GO metagenomics dataset^18^, resulting in an integrated diet-metagenomics dataset of 86 samples from 35 patients with 26.1 ± 1.2 million high-quality read pairs per sample (7.71 ± 0.36 gigabase pairs; **Table S8**). We correlated change in diet with change in gene family abundance. The 4 diet features with the largest number of significant diet-gene interactions were copper, carbohydrates, legumes, and diet quality (**Fig. 4a, Table S9**). Diet quality and carbohydrates were consistently altered during treatment (**Fig. 1b,f**), while copper and legumes were not (**Fig. 1g**).

**Figure 4:**
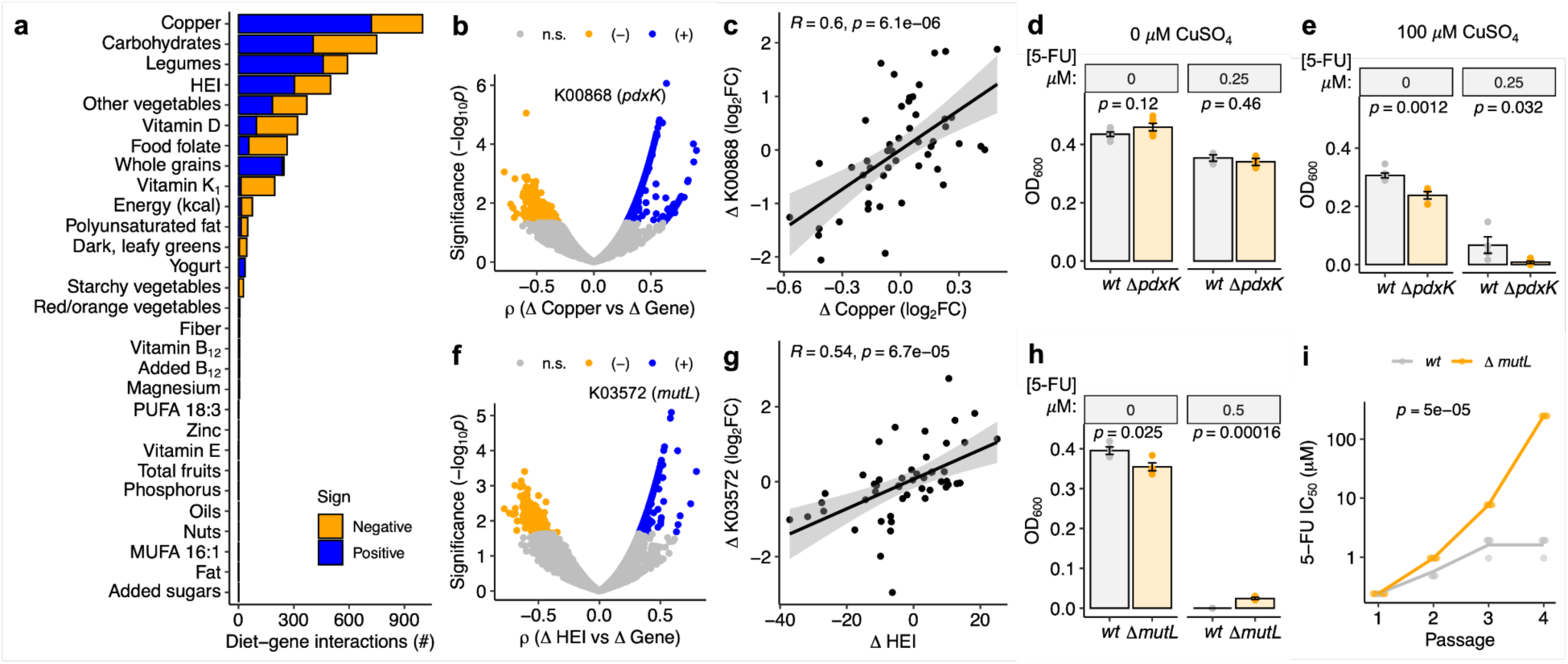
Dietary intake shifts correlate with micronutrient- and chemotherapy-related gut microbial genes. (**a**) Barplot of diet-gene interactions for each dietary variable. All diet variables with at least 1 significant diet-gene interaction are depicted. (**b**) Volcano plot of all copper-gene interactions. (**c**) Scatterplot of change in copper abundance vs change in pyridoxine kinase (K00868, *pdxK*) gene abundance in patients. (**d,e**) Carrying capacity (OD_600_) of *E. coli* BW25113 *wt* and Δ*pdxK* grown for 48 hours in M9-Glc ± 100 μM CuSO_4_ ± 0.25 μM 5-FU (*n*=4-6 replicates across 2 independent experiments). (**d**) *pdxK* deletion does not impact 5-FU toxicity. (**e**) *pdxK* deletion exacerbates copper toxicity in the presence or absence of 5-FU. (**f**) Volcano plot of all healthy eating index (HEI)-gene interactions. (**g**) Scatterplot of change in HEI vs change in mismatch repair (K03572, *mutL*) gene abundance in patients. (**h**) Carrying capacity (OD_600_) of *E. coli* BW25113 *wt* and Δ*mutL* grown for 48 hours in M9-Glc ± 0.5 μM 5-FU. *mutL* deletion partially rescues 5-FU toxicity (*n*=3 replicates). (**i**) 5-FU IC_50_ of *E. coli* BW25113 *wt* and Δ*mutL* serially passaged for 24 hours in M9-Glc ± varying concentrations of 5-FU as part of an experimental evolution experiment (see *Methods*). *mutL* deletion accelerates evolution of 5-FU resistance. *p*-values: (**a,b,f**) Spearman correlation of change in rank-transformed dietary intake vs change in rank-transformed central log ratio [CLR]-normalized KEGG ortholog gene family abundance, with Benjamini-Hochberg false discovery rate (FDR) correction across all tested genes for each dietary variable; (**c,g**) Linear regression of log₂ fold-change of dietary intake vs log₂ fold-change gene abundance between each pair of consecutive within-patient timepoints, with solid black line indicating the line of best fit and grey bands the 95% confidence interval; (**d,e,h**) one-way ANOVA; (**i**) Strain*passage interaction term from two-way ANOVA.

Because of the large quantity of copper-gene interactions, we opted to focus on copper. We identified 998 significant copper-gene interactions, including 721 positive and 277 negative interactions (**Fig. 4b, Table S10**). The top hit was an interaction between copper and pyridoxine kinase (K00868, *pdxK*), a gene that converts inactive vitamin B_6_ into its activated form of pyridoxal 5’-phosphate (PLP)^23^ (**Fig. 4b**). Increases in dietary copper were associated with increases in bacterial *pdxK* (**Fig. 4c**), with similar trends across all sub-cohorts (**Extended Data Fig. 7a-d**) and across taxonomically stratified *pdxK* abundances (**Extended Data Fig. 7e,f**). We turned to bacterial genetics to investigate the causal role of *pdxK* in bacterial drug and copper sensitivity. We grew *E. coli* BW25113 wild-type (*wt*) and Δ*pdxK*::*Kan^R^* in M9-Glc ± 250 nM 5-FU ± 100 μM CuSO_4_ (a Cu^2+^ source, matching the oxidation state of dietary copper^24,25^). Growth of the 2 strains was comparable in the presence and absence of 5-FU (**Fig. 4d**). However, CuSO_4_ significantly decreased the growth of Δ*pdxK*::*Kan^R^* relative to *wt* irrespective of 5-FU (**Fig. 4e**).

We next investigated diet quality-gene interactions given persistent decreases in HEI during treatment (**Fig. 1b**) and previously identified associations between HEI and microbiome composition (**Fig. 2c,e**). We identified 499 significant HEI-gene interactions, including 303 positive and 196 negative interactions (**Fig. 4f, Table S11**). The top hit was an interaction between HEI and a DNA mismatch repair gene (K03572, *mutL*), where increased dietary HEI was linked to increased bacterial *mutL* across all 3 sub-cohorts (**Fig. 4g, Extended Data Fig. 7g-j**) and across taxonomically stratified *mutL* abundances (**Extended Data Fig. 7k-l**). Higher baseline total *mutL* abundance was positively associated with hand-foot syndrome (HFS) development (**Extended Data Fig. 7m**), driven in part by *mutL* of unclassified origin (**Extended Data Fig. 7n**). In contrast, *Coprococcus*-derived *mutL* was negatively associated with HFS (**Extended Data Fig. 7o**), even though *Coprococcus* genus abundance was not (**Extended Data Fig. 7p**).

We explored how *mutL* impacted chemotoxicity, growing *E. coli* BW25113 wild-type (*wt*) and Δ*mutL*::*Kan^R^* in M9-Glc ± 500 nM 5-FU. While Δ*mutL*::*Kan^R^* growth was lower than *wt* in the absence of 5-FU, only Δ*mutL*::*Kan^R^* was able to grow in the presence of 5-FU (**Fig. 4h**). Next, we conducted experimental evolution of 5-FU resistance in *wt* and Δ*mutL*::*Kan^R^* strains. We evolved 8 lines (2 strains x 4 replicates) in M9-Glc medium with a 2-fold 5-FU dilution series. For 4 serial passages, cells from the highest 5-FU concentration with OD_600_ ≥ 0.03 were transferred to fresh medium at the same starting OD (**Fig. S2a**). In this setting, Δ*mutL*::*Kan^R^*developed 5-FU resistance significantly faster than *wt* (**Fig. 4i, Fig. S2b**). The MIC of the Δ*mutL*::*Kan^R^* strain increased 10,000-fold (**Fig. S2b**), consistent with tolerance rather than metabolism since constitutive expression of the 5-FU-metabolizing gene *preTA* improves MIC only 4-fold^19^.

We next examined whether any significant diet-associated microbial functions were related to pathways with previously reported relevance in CRC development or treatment response, focusing on butyrate^26,27^, bile acids^28^, aryl hydrocarbon receptor (AhR) ligands^29,30^, vitamin K_2_^11^, and drug metabolism^18,19^. Increasing copper intake was linked to increased butyrate production genes and pathways (**Extended Data Fig. 8a-d**), which in turn were associated with treatment efficacy (**Extended Data Fig. 8e-g**). Increased diet quality and vegetables correlated with increased bile salt hydrolase and *baiB* abundance (**Extended Data Fig. 8h-i**), while increases in dark leafy greens and vitamin K_1_ correlated with decreases in AhR ligand genes *tyrB/araT* and *tnaA* (**Extended Data Fig. 8k-m**). Vitamin K_2_ synthesis genes were linked to higher carbohydrate, fruit, and fiber intake (**Extended Data Fig. 8n**). No diet features were associated with variation in abundance of the *preTA* operon involved in 5-FU inactivation (FDR > 0.2 for all).

Finally, we investigated diet-*Fusobacterium* interactions given that *Fusobacterium nucleatum* is enriched in CRC patients^31^ and is associated with worse treatment outcomes^32,33^. The *Fusobacterium* genus was detected in <15% of samples by 16S rRNA gene sequencing and metagenomics, leading us to quantify *F. nucleatum* using a validated qPCR assay (**Fig. 5a, Extended Data Fig. 9a-c**). *F. nucleatum* was detected in 93% of samples by qPCR (**Fig. 5a**), including at least 1 sample from every patient (100% prevalence). Estimated abundances were significantly correlated across all 3 methods (**Extended Data Fig. 9d-f**), with qPCR abundances used for all subsequent analyses.

**Figure 5:**
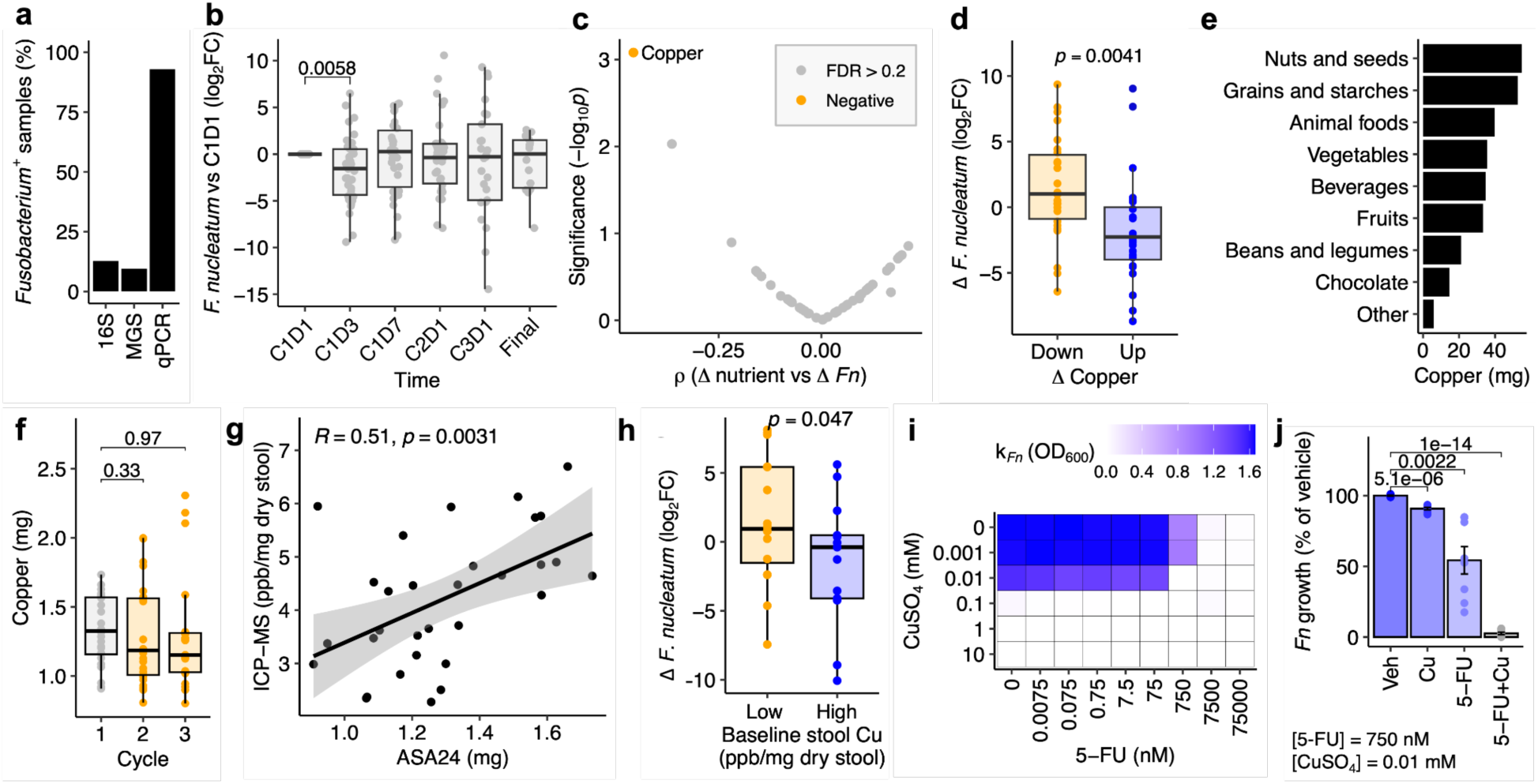
Copper suppresses *Fusobacterium nucleatum* in patients and *in vitro*. (**a**) Detection of *Fusobacterium* by 16S rRNA gene sequencing (16S), metagenomic sequencing (MGS), and quantitative PCR (qPCR). (**b**) Change in *F. nucleatum (Fn)* abundance during treatment, measured by qPCR. (**c**) Volcano plot of interactions between dietary micronutrients and *F. nucleatum* abundance. Each point represents a single micronutrient. (**d**) Change in *F. nucleatum* abundance vs change in copper intake. (**e**) Barplot of copper source. (**f**) Copper intake during treatment. (**g**) Correlation between C1 dietary copper intake (ASA24) and baseline stool copper (ICP-MS). (**h**) Change in *F. nucleatum* abundance during cycle 1 vs baseline stool copper levels (Low/High relative to median). (**i**) Carrying capacity of *F. nucleatum* (k*_Fn_*) grown for 24 hours in BHI^CHV^ ± CuSO_4_ (0, 0.001, 0.01, 0.1, 1, 10 mM) and 5-FU (0.0075, 0.075, 0.75, 7.5, 75, 750, 7500, 75000 nM; *n*=2-8 replicates pooled from 3 independent experiments). (**j**) *F. nucleatum* growth when treated with vehicle (Veh) ± 0.01 mM CuSO_4_ ± 750 nM 5-FU (*n*=8 replicates pooled from 3 independent experiments). *p*-values: (**b,j**) Welch’s *t*-test; (**c**) Benjamini-Hochberg false discovery rate (FDR)-adjusted Spearman correlation for change in rank-transformed dietary intake vs change in rank-transformed diversity between each pair of consecutive within-patient timepoints, with FDR<0.2 called as significant; (**d,f**) Mann-Whitney U test; (**g**) Pearson correlation; (**h**) one-sided Welch’s *t*-test.

Pre-treatment *F. nucleatum* abundance varied by orders of magnitude (**Extended Data Fig. 9g**). *F. nucleatum* abundance was significantly lower in patients with yoCRC (Mann-Whitney FDR < 0.2, **Extended Data Fig. 9h**). *F. nucleatum* did not vary with stage, tumor sidedness, BMI category, or sex (Mann-Whitney FDR > 0.2). High pre-treatment *F. nucleatum* was associated with worse response (**Extended Data Fig. 9i**).

*F. nucleatum* was significantly depleted during the first 3 days of chemotherapy (**Fig. 5b**), with no consistent differences later in treatment. *F. nucleatum* enrichment during treatment was associated with worse response (**Extended Data Fig. 9j**). To determine whether any diet-derived compounds might sensitize *F. nucleatum* to chemotherapy, we correlated changes in *F. nucleatum* with shifts in each of the 46 ASA24-quantified micronutrients. Increased copper intake was linked to *F. nucleatum* depletion (**Fig. 5c,d, Extended Data Fig. 9k-m**). This association was similar across sex, stage, tumor sidedness, BMI, and age (**Table S12**). Dietary copper was derived from a variety of sources (**Fig. 5e**) and was not significantly altered during treatment (**Fig. 5f**).

We validated these trends by using inductively coupled plasma mass spectrometry (ICP-MS) to directly measure ion levels in baseline stool samples (*Methods*, **Fig. S3a,b**). Of all measured ions, only stool copper was significantly correlated with dietary intake (**Fig. S3c**, **Fig. 5g**). Pre-treatment stool copper levels were associated with *F. nucleatum* suppression during chemotherapy (**Fig. 5h**), with similar trends observed across all subcohorts (**Fig. S3d**).

*In vitro* growth experiments supported a causal role of copper in suppressing *Fusobacterium nucleatum*. We grew *F. nucleatum* DSM15643 anaerobically in a gradient of CuSO_4_ and 5-FU (**Fig. 5i, S4**). *F. nucleatum* carrying capacity was slightly decreased by 10 µM copper and halved by 750 nM 5-FU (**Fig. 5i,j, S4**). A combination of 10 µM copper and 750 nM 5-FU completely inhibited *F. nucleatum* (**Fig. 5i,j, S4**), indicating a synergistic antimicrobial effect of copper and 5-FU for this pathogenic species.

Finally, to assess the *in vivo* relevance of dietary copper for *Fusobacteriota* in an animal model, we re-analyzed published 16S rRNA gene sequencing data^34^ from the gut microbiota of 25 turtles treated with varying copper concentrations (**Extended Data Fig. 10a,b**). Though overall microbial diversity was unaffected by copper (**Extended Data Fig. 10c,d)**, copper significantly suppressed *Fusobacteriota* levels in a dose-dependent manner (**Extended Data Fig. 10e**).

## DISCUSSION

Our results demonstrate that dietary intake markedly changes during oral fluoropyrimidine chemotherapy. We observed a decline in diet quality during treatment marked by increased refined grains and reduced nut consumption. We also identified multiple individual nutrients that declined during chemotherapy, including vitamin K_1_, vitamin E, and theobromine. Our findings extend prior observations of decreased vitamin D intake during CRC treatment^35,36^ to a wider array of fat-soluble micronutrients. In particular, vitamin E prevents chemotherapy-induced neuropathy and mucositis^37–39^, making it a compelling potential target for dietary intervention or supplementation.

Baseline sugar and theobromine correlated with treatment outcomes. Surprisingly, we found fewer dose adjustments in patients with higher baseline added sugar intake, despite current American Cancer Society guidelines that advise against all sugar-sweetened beverages^40^. Although higher BMI has been associated with chemoprotection^41^, BMI did not explain our added sugars-dose adjustment relationship. One potential mechanism is that lower-fat, higher-carbohydrate intake may decrease farnesoid x receptor activity^42^, resulting in increased expression of 5-FU catabolism enzyme dihydropyrimidine dehydrogenase and thus reduced toxic drug levels^43^. Interestingly, high baseline theobromine intake was associated with protection from hand-foot syndrome (HFS), often a dose-limiting toxicity of fluoropyrimidine treatments including CAP, in patients across independent patient cohorts in the USA and the Netherlands. Whether dietary theobromine prevents HFS merits further study, given that theobromine inhibits HFS-relevant TRPV1-mediated sensory activation^44,45^.

Our results also highlight the potential for diet and chemotherapy to act together to shape microbiome composition. Microbiome dynamics reflected shifts in overall diet quality and chocolate-derived theobromine, consistent with prior work that pure and chocolate-derived theobromine can directly alter gut microbiome composition in animal models^46^. Strikingly, diet shifts explained variation in one third (5/15) of chemotherapy-associated bacteria. This includes *F. prausnitzii*, which is elevated by vitamin K_1_-rich plant-based diets^47^ and has been linked to improved immunotherapy outcomes in mice and patients^48,49^. Though *F. prausnitzii* was consistently depleted during chemotherapy across our cohort, subjects who decreased their vitamin K_1_ intake experienced a 20-fold greater depletion, motivating further study of plant-based diets during combination chemo- and immunotherapy.

Many of the strongest diet-microbe interactions involved food-derived bacteria, illustrating one way stool can reflect host dietary intake. In this study, more patients submitted stool (*n*=110) than filled out diet surveys (*n*=86) at each paired timepoint, highlighting the value of diet estimation from stool. Future work should explicitly integrate food-derived bacterial abundance with other methods to infer diet from stool, such as meta-barcoding^50^, metaproteomics^51^, and metagenomic estimation of dietary intake^52^.

Notably, while the antimicrobial properties of copper surfaces are well characterized^53,54^, far less is known about the impact of dietary copper on the human gut microbiota. We identified hundreds of copper-microbiome interactions and validated the top relationship in cell assays: a positive interaction between copper and vitamin B_6_-activating pyridoxine kinase (*pdxK*). Strikingly, *pdxK* deletion exacerbated copper toxicity in *E. coli*. A possible mechanism is that vitamin B_6_ acts as a cofactor for synthesis of essential iron-sulfur clusters^55,56^, which are disrupted by copper^57,58^. Interestingly, *pdxK* knockdown increases copper sensitivity in mammalian cells^59^, raising the hypothesis that this vitamin B_6_-copper relationship is conserved across the tree of life.

Copper and 5-FU synergistically inhibited the clinically relevant bacterium *Fusobacterium nucleatum*. *F. nucleatum* uses the virulence factor FadA to bind to and invade epithelial cells^60^. FadA complexes with copper to drive toxic reactive oxygen species production^61^, potentially contributing to the observed copper sensitivity of *F. nucleatum*. Interestingly, *F. nucleatum* lacks *pdxK*^62^, which may further drive toxicity. Together, our findings in *E. coli* and *F. nucleatum* suggest dietary copper impacts multiple members of the human gut microbiota, extending prior work in environmental and swine-associated microbial communities^63–65^.

We were surprised that overall caloric intake did not significantly decline during chemotherapy. One explanation for this observation is that our diet surveys were administered during 3 off-treatment days immediately prior to the first dose of each cycle^66^, a study design decision made to limit patient burden during days of active treatment. Larger diet shifts may have occurred mid-cycle, consistent with our prior observations of more dramatic shifts in gut microbial gene abundance at cycle 1 day 7^18^. Future studies should assess diet during active treatment to determine if acute dietary changes occur and explain dramatic early-treatment microbiome shifts.

Though our findings raise several interesting hypotheses that were experimentally validated, our sample size was small (*n*=35); further validation of our results in larger cohorts is needed^67^. While many key findings were similar across sub-cohorts, cross-cohort variability was present and could reflect underlying biological differences or statistical noise (*n*<10 TAS102, CAP+IO cohorts). Assembling even our modest cohort required 4 years of recruitment across multiple clinical sites, underscoring the difficulty of longitudinal diet-microbiome studies during cancer treatment. Our limited efficacy and toxicity data were hypothesis-generating; future studies should include more granular toxicity assessment (*i.e.* Common Terminology Criteria for Adverse Events) and assessment of clinical outcomes (*i.e.* overall survival, progression-free survival, hospitalizations).

Overall, our findings provide support for the role of diet in shaping the gut microbiome during cancer treatment. Our results nominate numerous testable diet-microbiome-chemotherapy interactions for further exploration in mouse models and clinical studies, including using a vitamin K_1_-rich diet to enrich beneficial *F. prausnitzii* or a copper-rich diet to deplete cancer-driving *F. nucleatum*. Our approach highlights the utility of combining longitudinal diet analysis, longitudinal microbiome analysis, and cell assays, elevating diet from an overlooked confounder into a manipulable tool for future clinical microbiome studies.

## METHODS

### Study participants

As described previously^18^, the <u>G</u>ut microbiome and <u>O</u>ral fluoropyrimidines (GO) study was an observational clinical study registered at ClinicalTrials.gov (NCT04054908; **Extended Data Fig. 1a**). The GO study was approved by the UCSF Institutional Review Board, and all participants provided informed consent. Participants were recruited and sampled, without any compensation, at UCSF from the study start date (2018/04/13) to end date (2022/06/30). Inclusion criteria: (*i*) ≥18 years old; (*ii*) histologically confirmed colorectal adenocarcinoma; and (*iii*) expected to receive oral fluoropyrimidine therapy. Exclusion criteria: (*i*) HIV positive; (*ii*) chemotherapy, biologic, or immunotherapy in the previous 2 weeks; (*iii*) exposure to ≥2 weeks of antibiotics in the last 6 months; or (*iv*) exposure to antibiotics in the past 4 weeks. Patients were enrolled in 1 of 3 subcohorts based on their treatment plan: (A) oral capecitabine (CAP) as part of standard-of-care therapy (including concurrent rectal radiation therapy); (B) TAS-102 (trifluridine/tipiracil) ± Y-90 radioembolization (NCT02602327)^68^; (C) CAP and immunotherapy (pembrolizumab) (NCT03396926). CAP and TAS-102 were prescribed according to FDA labels and investigator discretion for oral tablet dosing. CAP tablets were taken twice daily on days 1–14 of a 21-day cycle (no RT) or on days of radiation only (RT), while TAS-102 tablets were taken twice daily on days 1–5 and 8–12 of a 28-day cycle.

### Dietary assessment

Participants were asked to complete up to 4 3-day dietary records using the National Cancer Institute’s Automated Self-Administered 24-hour Dietary Assessment Tool (ASA24) online system. Dietary information was collected during the 3 days immediately preceding the start of cycles 1-3 (C1-C3) and at the end of treatment (EOT)^69^ (**Fig. 1a**). 35 participants submitted at least 1 set of dietary records (**Table S1, Extended Data Fig. 1c**), while 25/35 submitted at least 2 timepoints of dietary records (**Extended Data Fig. 1c**). We opted for collection of 3-day dietary records for the following reasons: (1) 3-day dietary records can reliably capture usual dietary intake^70^, (2) intake immediately preceding stool collection was expected to be more strongly associated with the microbiome^4^, and (3) consecutive days reduced study burden on participants already managing the challenges of active cancer treatment. To minimize measurement error in diet data, we excluded record days at the extremes of caloric intake: >99^th^ percentile or <1^st^ percentile of caloric intake from NHANES data collected in 2017-2020^71,72^. In total, this resulted in the exclusion of 11 24-hour periods (10 records < 458 kcal, 1 record > 5105 kcal; **Extended Data Fig. 1b**) and the subsequent exclusion of 2 sample timepoints (Patient 6 Cycle 1 and Patient 11 Cycle 2). The valid days within each 3-day collection period were averaged to obtain mean daily nutrient intake for each patient for each sampling period, resulting in a dietary dataset of 86 timepoints from 35 patients (**Extended Data Fig. 1c,d** and **Table S1**). Healthy eating index (HEI)-2020 and alternative healthy eating index (AHEI) were calculated using custom R scripts adapted from publicly available SAS code from the National Cancer Institute^73^. Because the ASA24 nutrient output does not include *trans* fat, we excluded this category from calculation of the AHEI; as a result, the AHEI displayed here has a maximum value of 100 rather than 110. For individual nutrient and food group analysis, dietary variables were normalized for total energy intake using the residual method^74^. The residual method works as follows: given the diet intake D*_j_* and kcal C*_j_* for each patient *j*: (*i*) fit regression coefficients b_0_ and b_1_ for equation D = b_0_ + b_1_C across all patients simultaneously; (*ii*) compute the residuals r*_j_* = D*_j_* -(b_0_ + b_1_C*_j_*); and (*iii*) calculate the normalized D*_j,_*_adjusted_ = r_j_ + D_avg_.

### Recommended daily allowances

Recommended daily allowances were calculated for diet variables with daily intake recommendations in the USDA/HHS Dietary Guidelines for Americans (2025-2030) or the NIH Office of Dietary Supplements (2026) (**Table S4**). Sex-specific conservative thresholds were used to reduce the likelihood of classifying patients as deficient: for age-dependent recommendations, the lowest recommended intake threshold among adults was used, and for calorie-specific recommendations, a conservative reference intake of 1400 kcal/day was used (**Table S4**).

### Stool collection and sequencing

As described previously^18^, stool was collected from GO study participants at up to 7 timepoints spanning pre-to post-chemotherapy: baseline, cycle 1 day 1 (C1D1), cycle 1 day 3 (C1D3), cycle 1 day 7 (C1D7), cycle 2 day 1 (C2D1), cycle 3 day 1 (C3D1), and end-of-treatment (EOT) (**Fig. 1a; Tables S6, S8**). Dietary records were collected over the 3 days immediately preceding C1D1, C2D1, and C3D1 (denoted C1, C2, C3). 40 participants submitted at least 1 stool sample (**Table S1**), while 38/40 submitted at least 1 stool sample pre-treatment and at least 1 sample during treatment (**Tables S6, S8**). All participants who submitted dietary records also submitted a stool sample for the corresponding timepoint (**Tables S6, S8**).

Library preparation and sequencing were performed as described previously on all 222 collected stool samples from 40 CRC patients from the GO clinical study^18^ and on 156 samples from 56 CRC patients in an orthogonal Dutch clinical trial^11^ (**Table S13**). In the GO study, paired 16S rRNA gene sequencing (V4 region, primers 515F and 806R) and diet information was available for 85 samples from 35 patients (**Table S6**), while paired metagenomics and diet information was available for 86 samples from the same 35 patients (**Table S8**). Prior to downstream analysis, taxon and gene abundances were center log ratio (CLR)-transformed.

### 16S rRNA gene sequencing data processing

Primers and adapters were removed using the cutadapt trim-paired command in QIIME2 (v2020.11)^75^. Sequences were trimmed to 220 bp (forward) or 150 bp (reverse). Quality filtering, denoising, and chimera filtering were performed using dada2 (v1.18.0) with QIIME2 command denoise-paired^76^. Sequence length was filtered to 250-255 bp using QIIME2 command feature-table filter-seqs. Amplicon sequence variant (ASVs) taxonomy was assigned using the SILVA v138 database^77^. Sequence variants not present in at least 3 samples with at least 10 reads were removed. ASV relative abundances were then center log ratio (CLR)-transformed prior to downstream analysis.

### Metagenomics data processing

Demultiplexed reads underwent adapter trimming and quality filtering with FastP (v0.23.2)^78^, followed by host read removal via mapping to human genome (GRCh38) with BMTagger (v3.101)^79^. We annotated taxonomy with MetaPhlAn (v4.0.3)^80^. Genes were annotated with HUMAnN (v3.6). UniRef90 gene families were further mapped to KEGG Orthologous groups. Gene family abundances were CLR-transformed prior to downstream analysis.

### Metagenomic estimation of dietary intake

The metagenomic estimation of dietary intake tool (MEDI)^52^ was run with parameters read_length = 100, params.confidence = 0.1, params.threshold = 5 to estimate theobromine intake from stool on the metagenomics datasets from the GO study and Dutch study (**Table S13**). ASA24 and MEDI estimates of theobromine intake were significantly correlated in the GO study (ρ_Spearman_ = 0.4, *p* = 0.00079); ASA24 diet records were not conducted in the Dutch study.

### qPCR quantitation of Fusobacterium nucleatum

Genomic DNA (gDNA) was extracted using ZymoBIOMICs 96 MagBead DNA Kit (Zymo D4308) with 2 rounds of 5 min bead beating, as described previously^18^. gDNA was used to quantify *Fusobacterium nucleatum* using a previously published and validated primer pair (Fn1_FWD & Fn1_REV; **Table S14**) that targets the *F. nucleatum* 16S rRNA gene^81^ and a separate previously published and validated primer pair (16S_general_FWD & 16S_general_REV; **Table S14**) that targets general bacterial 16S rRNA genes^82^. Reactions were run using the following volumes: 4 μL of diluted gDNA (1:10 in deionized water), 1 μL of 3 μM primer stock, and 5 μL SYBR Select Master Mix for CFX (Thermo Fisher Scientific). Cycling conditions: 50°C for 2 min, 95°C for 2 min, and 40 cycles of 95°C for 15 sec and 60°C for 1 min^81,82^. Standard curves were generated with *F. nucleatum* DSM15643 gDNA from pure culture. *F. nucleatum* abundance was normalized to total 16S rRNA gene amplicons, as described^83^. Samples with *F. nucleatum* below the limit of detection (Ct > 40) were called *F. nucleatum*-negative. *F. nucleatum* was detected by qPCR in at least 1 sample for each patient (100% prevalence). For quantitative analyses, the abundance of *F. nucleatum*-negative samples was set to the minimum detected positive proportion.

### Fusobacterium nucleatum growth assays

All work with *F. nucleatum* was performed in an anaerobic chamber (Coy Laboratory Products) with an atmosphere of <4% H_2_, 20% CO_2_, and remainder N_2_. *F. nucleatum* DSM15643 was streaked on brain heart infusion (BHI) supplemented with L-Cysteine (0.05% w/v; Sigma W326305-100G), hemin (5 μg/ml), and vitamin K_3_ (menadione; 1 μg/ml), referred to as BHI^CHV^. Colonies were subcultured overnight in BHI^CHV^ at 37°C. 5-fluorouracil (5-FU; Sigma F6627) was dissolved directly into BHI^CHV^, filter sterilized, and assayed at 75,000, 7,500, 750, 75, 7.5, 0.75, 0.075, 0.0075, and 0 nM (Experiments 1-3, **Fig. 5i,j**) or 6,000, 3,000, 1,500, 750, 375, 187.5, 93.75 nM (Experiments 4-5, **Fig. S4**). CuSO_4_ (Sigma C1297-100G) was dissolved in sterile water to make a concentrated stock, filter sterilized, and assayed at 10,000, 1,000, 100, 10, 1, and 0 μM (Experiments 1-3, **Fig. 5i,j**) or 100, 10, 1, and 0 μM (Experiments 4-5, **Fig. S4**). A total of 5 µL seed culture diluted to OD_600_ = 0.1 was inoculated with 200 µL BHI^CHV^ media ± drug ± CuSO_4_ in a 96-well plate. Plates were covered with Breathe-Easy membranes (Sigma Z380059) to prevent evaporation and incubated anaerobically at 37°C for 24 h. OD_600_ readings were performed at the endpoint (BioTek EON plate reader), with background subtraction of sterile control wells. For Experiments 4-5 (**Fig. S4**), endpoint cultures were subjected to 10-fold serial dilutions in BHI^CHV^ (undiluted, 1:10, 1:10^2^, 1:10^3^, 1:10^4^, 1:10^5^, 1:10^6^), followed by plating 2.5 µL aliquots of each dilution onto BHI^CHV^ and anaerobic incubation at 37°C for 48 h for assessment of colony forming units (CFU) to corroborate OD results. The limit of detection (LOD) was estimated as 400 CFU/mL (recovery of a single colony from the undiluted sample volume of 0.0025 mL), with values below the LOD set to the LOD prior to analysis.

### Faecalibacterium prausnitzii growth assays

All work with *F. prausnitzii* was performed in an anaerobic chamber (Coy Laboratory Products) with an atmosphere of <4% H_2_, 20% CO_2_, and remainder N_2_. Healthy donor-derived *F. prausnitzii* WB-STR-0032 was obtained from Pendulum Therapeutics and streaked on BHI^CHV^ agar. Colonies were subcultured overnight in liquid BHI^CHV^ at 37°C. 5-FU was dissolved directly into BHI^CHV^, filter sterilized, and assayed at 0, 37.5, 75, and 150 μM. Sterile vitamin K_1_ (ThermoScientific L10575.06) was diluted to 1 mg/mL in sterile DMSO followed by further dilution in sterile deionized water and assayed at 0 and 3 µM. A total of 2 µL seed culture diluted to OD_600_ = 0.005 was inoculated with 200 µL media ± 5-FU ± vitamin K_1_ in a 96-well plate. Plates were covered and incubated anaerobically at 37°C for 24 h in a plate reader, with 1 min linear shake prior to OD_600_ readings every 15 min.

### Escherichia coli growth assays

*E. coli* BW25113 wild-type (*wt*), Δ*mutL*::KanR, and Δ*pdxK*::KanR were obtained from the Keio collection^84^ and streaked on Luria Broth (LB) agar with 30 µg/mL kanamycin. Colonies were subcultured overnight in modified M9 minimal media (Na₂HPO₄ 33.9 g/L, KH₂PO₄ 15.0 g/L, NaCl 2.5 g/L, NH₄Cl 5.0 g/L, 0.24 g/L MgSO₄, 0.011 g/L CaCl_2_) + 0.4% glucose (M9-Glc) at 37°C. 5-FU was dissolved directly into M9-Glc, filter sterilized, and assayed at 0, 250, and 500 nM. CuSO_4_ was dissolved into sterile water to make a concentrated stock, filter sterilized, and assayed at 0 and 100 µM. A total of 5 µL seed culture diluted to OD_600_ = 0.1 was inoculated with 100 µL M9-Glc ± drug (or ± copper for Δ*pdxK* experiments) in a 96-well plate. Plates were covered and incubated aerobically at 37°C for 48 h in a plate reader, with 1 min linear shake prior to OD_600_ readings every 15 min. We adapted an established protocol for experimental evolution of antibiotic resistance to study evolution of 5-FU resistance in Δ*mutL* vs wild-type^85^ (**Fig. S2**). Briefly, at each passage, bacteria were cultured in 100 µL M9-Glc ± varying concentrations of 5-FU in a 96-well plate. During the experiment, each culture line was exposed to 6 concentrations of antibiotics, corresponding to 5 wells with a 2-fold dilution series and 1 drug-free well. Bacteria were inoculated into these wells at an initial OD_600_ of 0.005. After 24 h growth at 37°C with 1 min shaking every 15 minutes, cells were sampled from the well with the highest drug concentration among the wells that had OD_600_ > 0.03, with sampled cells diluted to initial OD_600_ of 0.005 and inoculated into fresh medium in a new plate. 6 independent culture lines were maintained for each strain, with serial transfer conducted for 4 passages. IC_50_ was defined as the 5-FU concentration at which bacteria displayed 50% OD_600_ at 24 hours relative to that strain’s growth in drug free media, while MIC was defined as the minimum concentration at which there was no detectable bacterial growth (OD_600_ < 0.03).

### Inductively Coupled Plasma Mass Spectrometry (ICP-MS)

ICP-MS was used to quantify concentrations of copper, potassium, sulfur, sodium, phosphorus, magnesium, iron, calcium (^40^Ca and ^43^Ca), selenium, and zinc in stool. Briefly, baseline (BL) stool samples with >100 mg wet stool remaining were dried by lyophilization with pressure <100 mTorr for 72 hr in metal-free 15 mL conical tubes (Avantor 10026-322). Samples were then digested in 200 µL 30% H_2_O_2_ and 2 mL 70% Optima-grade nitric acid at 65°C overnight, then diluted with UltraPure water to 20% nitric acid for analysis. Elemental quantification was conducted using an Agilent 7700 ICP-MS attached to an ASX-560 autosampler. The settings for analysis were cell entrance = −40 V, cell exit = −60 V, plate bias = −60 V, OctP bias = −18 V, and helium flow = 4.5 ml/min. Optimal voltages for extract 2, omega bias, omega lens, OctP RF, and deflect were empirically determined. Calibration curves for elements were generated using ARISTAR ICP standard mix. Samples were introduced by a peristaltic pump with 0.5-mm-internal-diameter tubing through a borosilicate glass nebulizer (MicroMist). They were initially taken up at 0.5 RPS for 30 seconds, followed by 30 seconds at 0.1 RPS to stabilize the signal. Spectrum mode analysis was performed at 0.1 RPS, collecting 3 points across each peak and conducting 3 replicates of 100 sweeps for each element. The sampling probe and tubing were rinsed with 2% nitric acid for 30 seconds at 0.5 RPS between each sample. Data were acquired and analyzed using Agilent MassHunter (vA.01.02). For all ions, the signal-to-noise ratio exceeded 10:1, and standard curve R^2^ > 0.99 (copper: **Fig. S3**). Change in Fn abundance during cycle 1 was defined as C1D7 abundance (latest timepoint during cycle 1) minus BL abundance.

### Effect of copper supplementation on gut *Fusobacteriota* in vivo

A PubMed search for (“copper”) AND (“gut microbiota” OR “gut microbiome”) AND (“Fusobact*”) was performed on June 26, 2026, resulting in 9 studies: PMIDs 33311635, 39827614, 32036523, 41497648, 40706943, 41290094, 32064276, 42349178, 41065383. Of these, 5 were interventional animal studies (39827614, 32036523, 40706943, 41290094, 42349178); all but 40706943 reported decreased *Fusobacteriota* with copper supplementation. Sequencing data was publicly available only for PMID 39827614 (Ali *et al.*, 2025^34^).

Paired-end 16S rRNA amplicon sequencing reads from PMID 39827614 were downloaded via SRA Toolkit and processed using the Nephele QC pipeline v0.2.1^86^ with default parameters (Trimmomatic v0.39^87^; BBMerge v39.38^88^; FastQC v0.12.1^89^; MultiQC v1.33^90^). Quality-trimmed paired-end reads were then analyzed using the Nephele QIIME2 pipeline v0.1.6^75,86^ with default parameters (closed-reference clustering at 97% identity; trim length 200bp; QIIME2 2024.09 backbone full-length reference database; scikit-learn^91^ classifier trained on reference database). All samples contained >10,000 high-quality reads. Sequence variants not present in at least 2 samples with at least 5 reads were removed. Copper-treated samples from Day 14 and Day 28 were grouped for analysis.

### Modeling and statistical analysis

#### Software

R (v4.4.2) was used for all analysis and visualization^92^. The lme function from the nlme package (v3.1.167) was used for mixed-effects modeling^93^. The cor.test, t.test, and wilcox.test function from stats (v4.4.2) were used for Pearson correlations, Spearman correlations, Welch’s *t*-tests, Wilcoxon signed-rank tests, Mann-Whitney U tests. Packages ggplot2 (v.3.5.2) and ggpubr (v0.6.1) were used for plotting^94,95^.

#### Regimen heterogeneity

We pooled all patients across all subcohorts to increase statistical power. For key findings, we examined trends stratified by cohort (*i.e.* **Extended Data Fig. 2k**, **4l, 5g-i, 7a-d,g-j; Fig. S3d**), though small sample size limited our ability to perform robust sub-cohort statistical analysis (*n*=6 TAS102, *n*=9 CAP+IO).

#### Multiple hypothesis testing

For all statistical testing, whenever multiple tests were performed, Benjamini-Hochberg false discovery rate (FDR) correction was applied using the p.adjust function, with FDR < 0.2 called as significant.

#### Diet variable selection

Pearson correlations between all dietary variables from ASA24 were computed, with the less general variable of each highly correlated pair (ρ > 0.85) removed prior to analysis to enhance power (**Fig. S1a, Table S2**). When more than 2 variables were all highly correlated, we kept only 1 variable from each cluster, ensuring that ρ > 0.85 with all removed variables in the cluster (**Fig. S1a, Table S2**). For example, CHOLE was highly correlated with both CHOLN and P204 (ρ > 0.85), leading us to keep CHOLE and drop CHOLN and P204 (**Fig. S1a, Table S2**). Food groups with 0 consumption in >60% of the timepoints were also removed (**Table S2**), resulting in 79 ASA24 dietary features used for analysis (including HEI; **Table S3**).

#### Diet vs time

For modeling diet variables vs time, we filtered to only include patients with at least 2 timepoints. To take advantage of the longitudinal nature of the dataset, we used a mixed effects model of Diet ∼ PrePost + 1|Pt, *i.e.* all post-cycle 1 timepoints from a patient were assigned to the same “post” group. All significant hits (FDR<0.2) were plotted with boxplots, with intake at cycle 1 compared to intake at cycle 2 and at cycle 3 using the Wilcoxon signed-rank test. End-of-treatment was not plotted since only 8 patients submitted a dietary record at this timepoint, and the date of the end-of-treatment timepoint relative to cycle 1 varied by months from patient to patient^18^.

#### Diet and clinical variables vs outcome

For modeling change in diet vs outcomes, we used a Welch’s *t*-test to compare change in each diet feature between cycle 2 and cycle 1 to each outcome. For modeling baseline diet and clinical variables vs outcome, we used the Welch’s *t*-test for diet variables and the Fisher exact test for clinical variables. Top hits were plotted as boxplots, with intake from each outcome group compared using a Welch’s *t*-test. “BMI category” was binarized as BMI above vs below 25, yoCRC was binarized as age <50 at diagnosis, tumor sidedness was binarized as left vs right (distal vs proximal to splenic flexure), and stage was binarized as stage IV vs stage I-III. To assess whether sex, BMI category, yoCRC, tumor sidedness, or stage confounded the association between baseline diet and treatment outcomes, we compared odds ratios from logistic regression models with and without each metadata variable, with a >10% change of odds ratio towards 1 defined as evidence of effect attenuation.

#### Microbiome diversity calculations

Alpha diversity (observed) was calculated as the sum of observed genera. Alpha diversity (Shannon) was calculated by applying the diversity function from vegan (v2.7.1) to genus counts^96^. Beta diversity was calculated by applying the dist function to CLR-transformed genera composition (CLR-Euclidean ordination). PERMANOVA testing comparing baseline microbiome composition to GI toxicities was performed with the adonis2 function from the vegan package.

#### Modeling change in diet vs change in microbiome

We used Spearman correlation to assess the relationship between change in dietary intake and change in microbiome variables. To do this, we utilized every pair of consecutive within-patient timepoints with at least 1 nonzero value for diet and for the microbiome variable (*i.e.* a pair was excluded if there was no change in diet or no change in the microbiome feature); if fewer than 10 pairs fulfilled this criteria for a given diet-microbiome combination, that interaction was not tested. In our prior analysis, we identified 43 ASVs as differentially abundant over time^18^; only 15 of these ASVs were present in ≥10 pairs and evaluable for diet analysis (**Table S7**). Change in microbiome was calculated as the difference in rank-transformed number of observed genera (alpha diversity), Shannon index (alpha diversity), CLR-Euclidean distance (beta diversity), CLR-transformed ASV abundance (taxa), or CLR-transformed KEGG ortholog abundance (genes). To increase power and decrease the potential impact of contamination, we only tested taxa/genes that were detected in at least 25 samples (10% prevalence filter)^97^. Change in dietary intake was computed as the difference in rank-transformed dietary intake. For comparisons with beta diversity, the absolute value of the change in rank-transformed dietary intake was used (*i.e.* large increases or decreases in microbiome-relevant dietary features are expected to increase the distance in CLR-Euclidean ordination space).

#### Visualization of change in diet vs change in microbiome

Pearson correlation was used to visualize the relationship between change in dietary intake and change in microbiome since Spearman correlation cannot produce a line of best fit, with geom_smooth(method = “lm”) used to plot a linear regression line of best fit and 95% confidence interval. Each data point represents a pair of consecutive within-patient timepoints with at least 1 nonzero value for diet and for the microbiome variable (*i.e.* data points with no change in diet or no change in microbiome were excluded). Log_2_ fold change was computed between each feature at time *t* and *t*+1 within the given pair, with a pseudocount equal to the feature’s minimum positive value in the dataset added prior to log transformation. Boxplots were made by binning diet changes into increasing or decreasing intake, with the Mann-Whitney U test used for statistical comparison. For visualization of ASV dynamics over time, patients were binned based on whether their intake of a diet feature increased or decreased between cycle 1 and a single later cycle. The mean ± standard error of ASV abundance relative to baseline for patients in each of these groups was plotted across all treatment timepoints, including timepoints with no diet information available, with two-way analysis of variance (ANOVA) used to test group differences.

#### Overrepresentation of food-derived bacteria

*Streptococcus and Lactobacillus* spp. represent prevalent and abundant food-derived bacteria^22^. Fisher’s exact test was used to test representation of these ASVs in diet-ASV interactions.

#### Fusobacterium nucleatum qPCR analysis

Because 16S rRNA gene sequencing of the V4 region classifies most ASVs at the genus but not species level^98^, we opted to compare *Fusobacterium* abundance at the genus level from 16S and MGS vs *F. nucleatum* species level abundance by qPCR. Pearson correlation was used to test standard curves and correlations between different quantification methods. Binarized patient metadata (BMI, sex, age, stage, tumor sidedness) were compared with pre-treatment (C1D1) *F. nucleatum* abundance via one-sided Mann-Whitney U tests, with significance called when both *p* < 0.05 and FDR < 0.2. We assessed the relationship between change in intake of 46 micronutrients and change in *F. nucleatum* using Spearman correlation. We used every pair of consecutive within-patient timepoints with at least 1 nonzero value for diet. Changes in diet and *F. nucleatum* were calculated as the difference in rank-transformed intake and proportion, respectively. Boxplots were made by binning diet changes into increasing or decreasing intake, with Mann-Whitney U test used for statistical comparison of *F. nucleatum* proportion.

#### In vitro experiments

Growth was compared using one-way ANOVA (equal variances) or Welch’s *t*-test (unequal variances). We tested IC_50_ changes in the experimental evolution study via two-way ANOVA (strain*passage interaction term).

## DATA AVAILABILITY

16S rRNA and metagenomic gene sequencing data used in this study are publicly available in the Sequence Read Archive (SRA) under accession number PRJNA1169175. All data is available on Github (https://github.com/turnbaughlab/2025_Trepka_GODiet); tables with clinical metadata and 3-day averaged ASA24 raw and residual-normalized data are in subdirectory DietAndClinicalData (https://github.com/turnbaughlab/2025_Trepka_GODiet/tree/main/!DietAndClinicalData). All other data is available from the corresponding author upon request.

## CODE AVAILABILITY

All code is publicly available on GitHub (https://github.com/turnbaughlab/2025_Trepka_GODiet).

## ACKNOWLEDGEMENTS

Funding was provided by the National Institutes of Health (R01CA255116, R01DK114034, and R01HL122593 to P.J.T.; F30CA257378 to TSK) and the Parkinson’s Disease Foundation (PF-LAUNCH-1244609 to CAO). P.J.T is a Biohub, San Francisco, Investigator. G.P. is supported by the President’s Postdoctoral Fellowship Program (PPFP). Sequencing was performed at Biohub, San Francisco and the UCSF Center for Advanced Technology. Diagrams were created with <u>Biorender.com</u>. *Faecalibacterium prausnitzii* WB-STR-0032 was provided by Pendulum Therapeutics.

## EXTENDED DATA FIGURES AND FIGURE LEGENDS

**Extended Data Figure 1:**
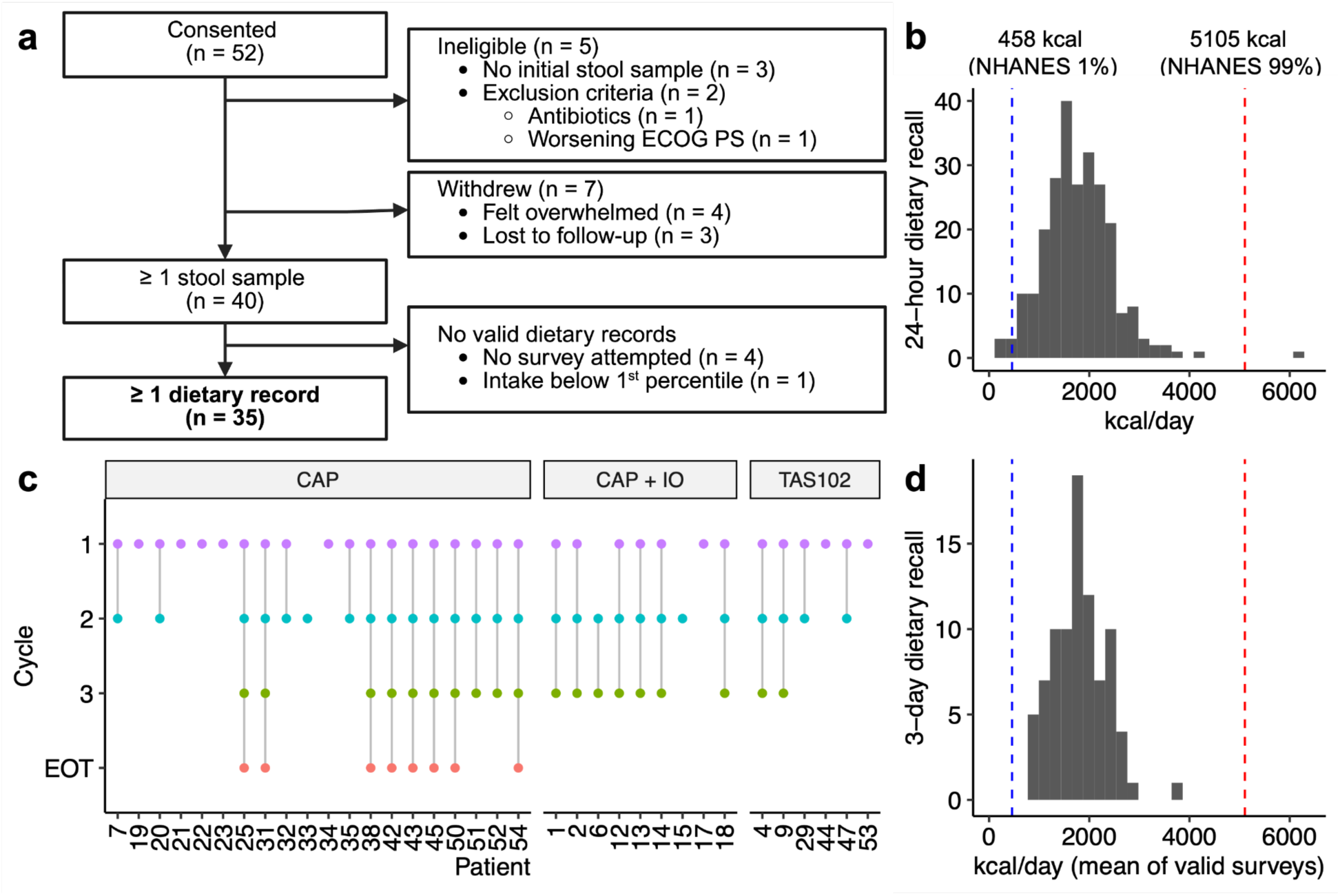
GO diet study design. (**a**) CONSORT diagram. (**b**) Energy intake distribution for each individual 24-hour record. 24-hour records below the 1st percentile or above the 99th percentile of National Health and Nutrition Examination Survey (NHANES) intake were excluded. (**c**) Distribution of included samples (diet and stool sequencing data available). (**d**) Average daily energy intake distribution over 3-day dietary record for samples from (**c**).

**Extended Data Figure 2:**
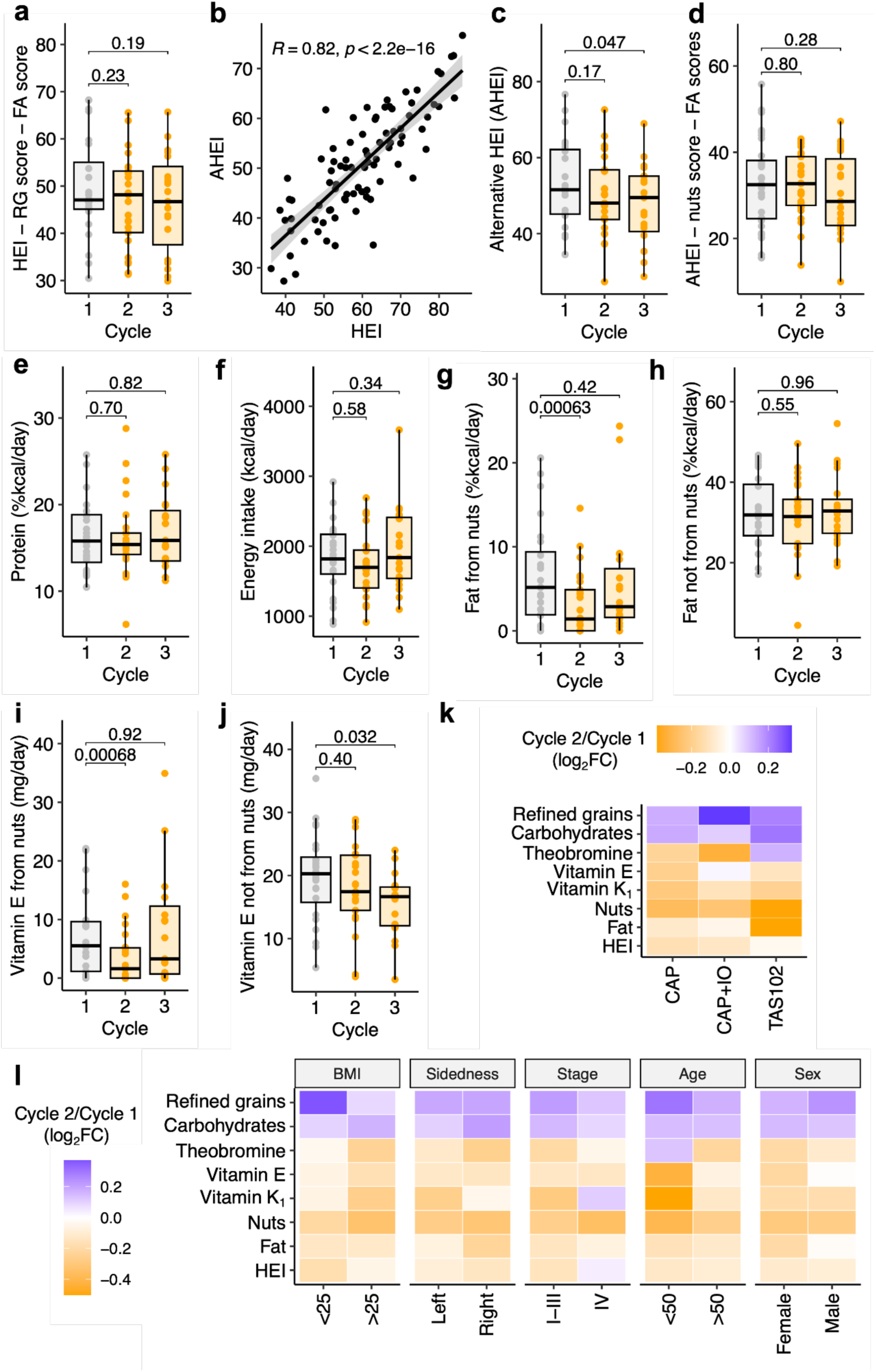
Dietary intake during chemotherapy. (**a**) HEI excluding the refined grains (RG) score and fatty acids (FA) score was unchanged during the study. (**b**) Correlation of HEI and Alternative HEI (AHEI). (**c**) AHEI decreased during the study. (**d**) AHEI excluding the nuts score and fatty acids scores was unchanged during the study. (**e,f**) Change in protein (**e**) and energy intake (**f**) during the study. (**g,h**) Change in nut-derived fat (**g**) and all other fat (**h**) during the study. (**i,j**) Change in nut-derived vitamin E (**i**) and all other vitamin E (**j**) during the study. (**k-l**) Changes in dietary intake were similar across all 3 subcohorts (**k**) and across patient metadata categories of BMI, tumor sidedness, tumor stage, age/yoCRC, or sex (**l**), as measured by mean log_2_ fold change (log_2_FC). Purple (orange) represents an increase (decrease) in intake from C1 to C2. *p*-values: Wilcoxon signed-rank test (**a,c-j**), Pearson correlation with solid black line indicating the line of best fit and grey bands the 95% confidence interval (**b**).

**Extended Data Figure 3:**
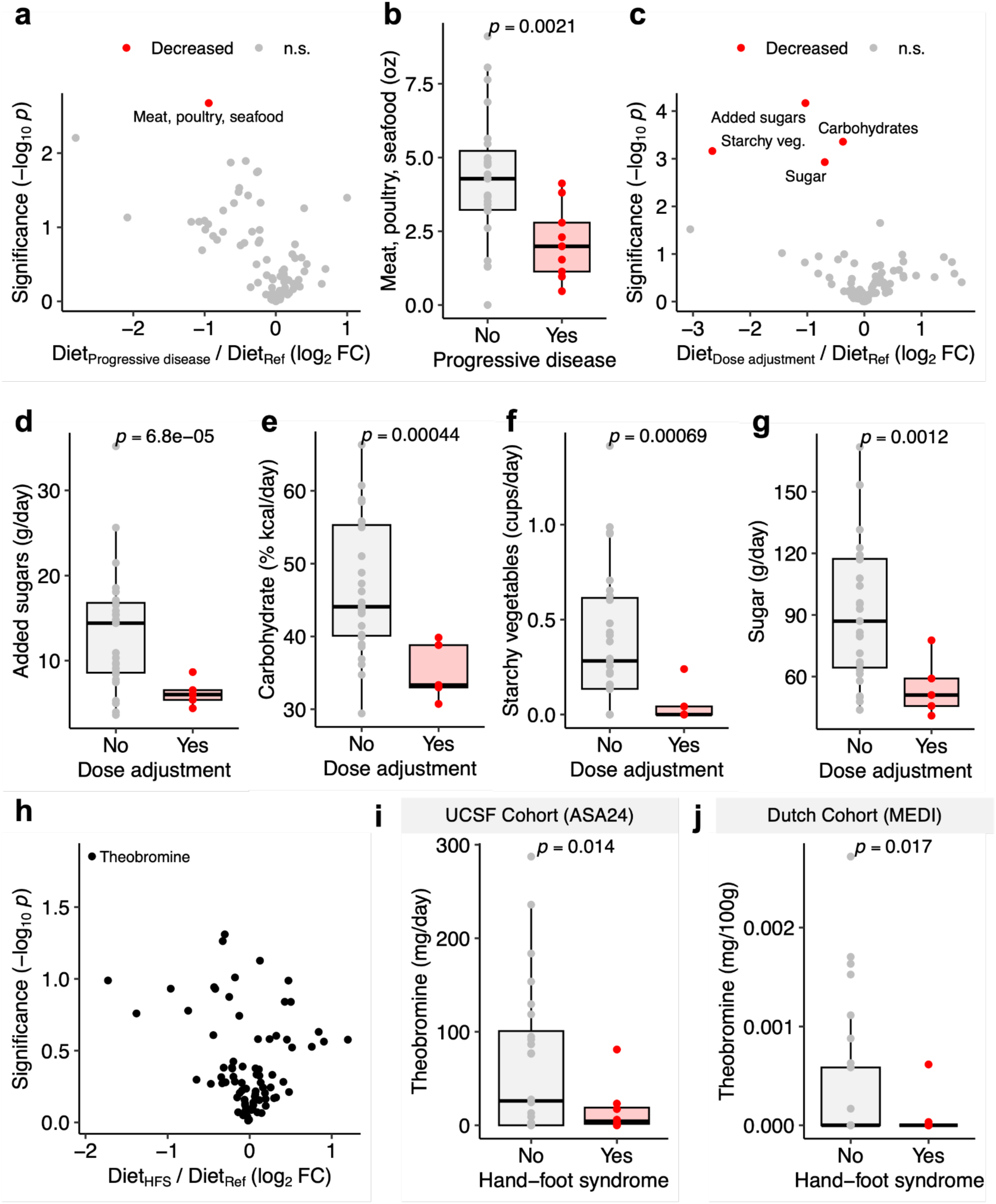
Baseline dietary intake is associated with drug efficacy and toxicity. (**a**) Volcano plot of baseline (C1) dietary intake in patients who had progressive disease vs stable disease or response. Each point represents a dietary variable. (**b**) Baseline meat, poultry, and seafood intake vs progressive disease status. (**c**) Volcano plot of baseline dietary intake in patients who required dose adjustments (delay or reduction) vs no dose adjustments. Each point represents a dietary variable. (**d-g**) Dose adjustment status vs baseline intake of added sugars (**d**), carbohydrate (**e**), starchy vegetables (**f**), and sugar (**g**). (**h**) Volcano plot of baseline dietary intake in patients who experience hand-foot syndrome (HFS) vs no HFS. Each point represents a dietary variable. (**i**) Hand-foot syndrome vs ASA24-quantified baseline theobromine intake in the GO cohort. (**j**) Hand-foot syndrome vs MEDI^52^-estimated baseline theobromine intake in an independent Dutch cohort^11^. *p*-values: Welch’s *t*-test for all. Benjamini-Hochberg false discovery rate correction was performed for (**a,c**), with FDR<0.2 called as significant.

**Extended Data Figure 4:**
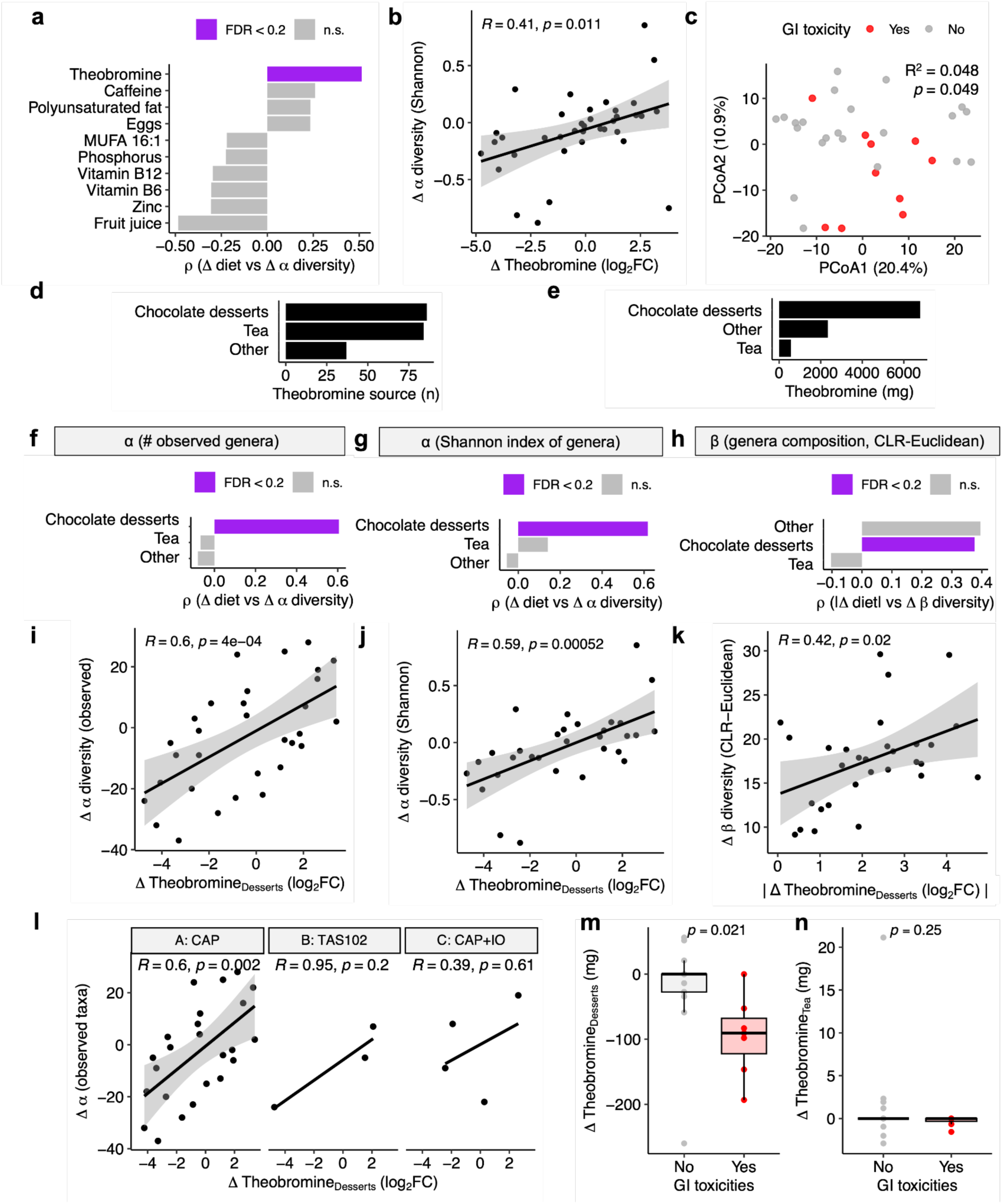
Dessert-derived theobromine intake shifts correlate with altered gut microbiome diversity. (**a**) Top 10 diet variables associated with shifts in alpha diversity (Shannon index of genera composition). (**b**) Change in Shannon index of genera composition vs change in theobromine intake. (**c**) Baseline microbiota composition is significantly associated with development of gastrointestinal toxicities. (**d,e**) Number of survey items (**d**) and total theobromine intake in mg (**e**) by food source. (**f-h**) Source-stratified theobromine vs change in alpha diversity (number of observed genera) (**f**), alpha diversity (Shannon index of genera) (**g**), and beta diversity (CLR-Euclidean distance on genera composition) (**h**). (**i-k**) Change in chocolate dessert-derived theobromine vs change in alpha diversity (number of observed genera) (**i**), alpha diversity (Shannon index of genera) (**j**), and beta diversity (**k**). **(l)** Chocolate dessert-derived theobromine-alpha diversity relationships were similar across all 3 subcohorts. (**m,n**) Gastrointestinal (GI) toxicities vs change in theobromine derived from chocolate desserts (**m**) or tea (**n**). *p*-values: (**a,f-h**) Benjamini-Hochberg false discovery rate (FDR)-adjusted Spearman correlation for change in rank-transformed dietary intake vs change in rank-transformed diversity [signed for (**a,f,g**), absolute value for (**h**)] between each pair of consecutive within-patient timepoints, with FDR < 0.2 called as significant; (**b,i-l**) Linear regression of log₂ fold-change of dietary intake [absolute value for (**k**)] vs log₂ fold-change of diversity between each pair of consecutive within-patient timepoints, with solid black line indicating the line of best fit and grey bands the 95% confidence interval; (**c**) PERMANOVA test; (**m-n**) Welch’s *t*-test.

**Extended Data Figure 5:**
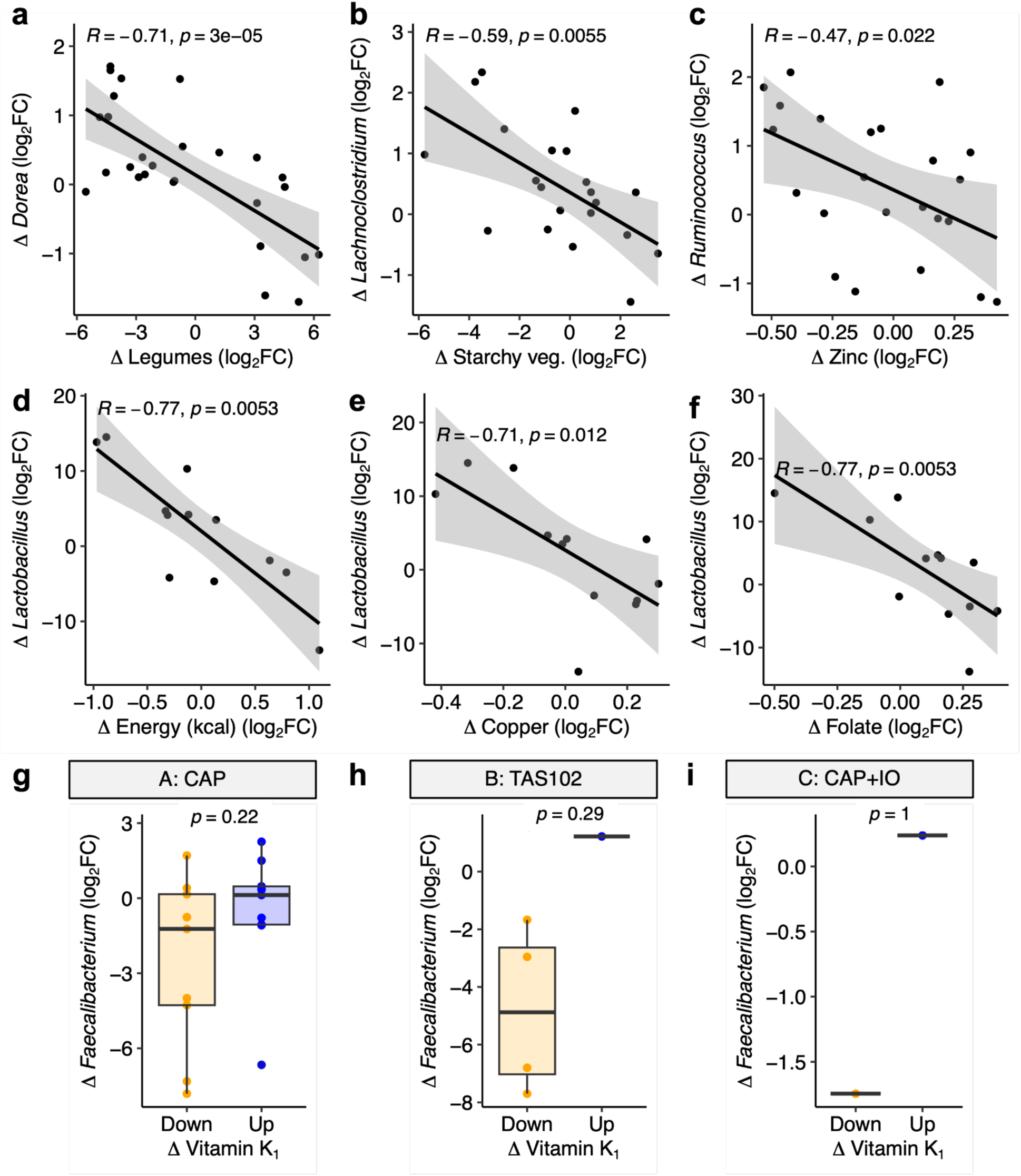
Shifts in diverse food groups and micronutrients correlate with shifts in CAP-altered gut bacteria. (**a-f**) Change in ASV abundance vs change in dietary intake for the 6 significant negative interactions from Fig. 3a and **Table S7**: legumes-*Dorea* (ASV9) (**a**), starchy vegetables-*Lachnoclostridium* (ASV3) (**b**), zinc-*Ruminococcus* (ASV13) (**c**), and energy-, copper-, and folate-*Lactobacillus* (ASV4) (**d-f**). (**g-i**) Change in *Faecalibacterium* ASV abundance vs change in vitamin K_1_ intake stratified by sub-cohort: CAP (**g**), TAS102 (**h**), CAP+IO (**i**). Each point represents the difference between 2 consecutive timepoints within a single patient. *p*-value: (**a-f**) linear regression of log₂ fold-change of dietary intake vs log₂ fold-change ASV abundance between each pair of consecutive within-patient timepoints, with solid black line indicating the line of best fit and grey bands the 95% confidence interval; (**g-i**) Mann-Whitney U test.

**Extended Data Figure 6:**
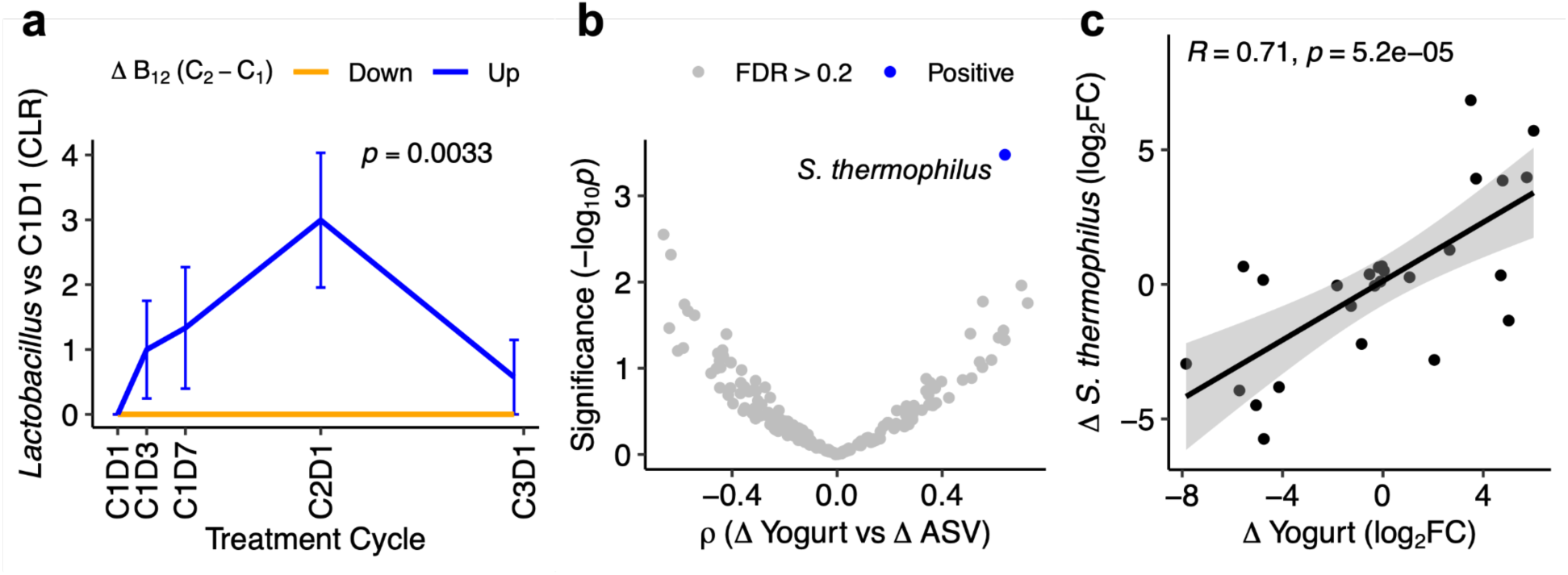
Variation in intake of probiotic foods explains variation in abundance of food-derived bacteria. **(a)** Change in *Lactobacillus* ASV abundance stratified by the change in vitamin B_12_ intake between C1 and C2. *p*-value: Group (diet) term from two-way ANOVA. **(b,c)** Variation in yogurt intake explains variation in abundance of yogurt-associated *Streptococcus thermophilus*. (**b**) Volcano plot of yogurt-ASV interactions. *p*-value: Spearman correlation of change in rank-transformed dietary intake vs change in rank-transformed central log ratio (CLR)-normalized ASV abundance. Benjamini-Hochberg false discovery rate (FDR) correction was applied to all *p*-values, with FDR<0.2 called as significant. (**c**) Change in yogurt vs change in *S. thermophilus* abundance between each pair of consecutive within-patient timepoints*. p*-value: Linear regression of log₂ fold-change of yogurt intake vs log₂ fold-change *S. thermophilus* abundance, with solid black line indicating the line of best fit and grey bands the 95% confidence interval.

**Extended Data Figure 7:**
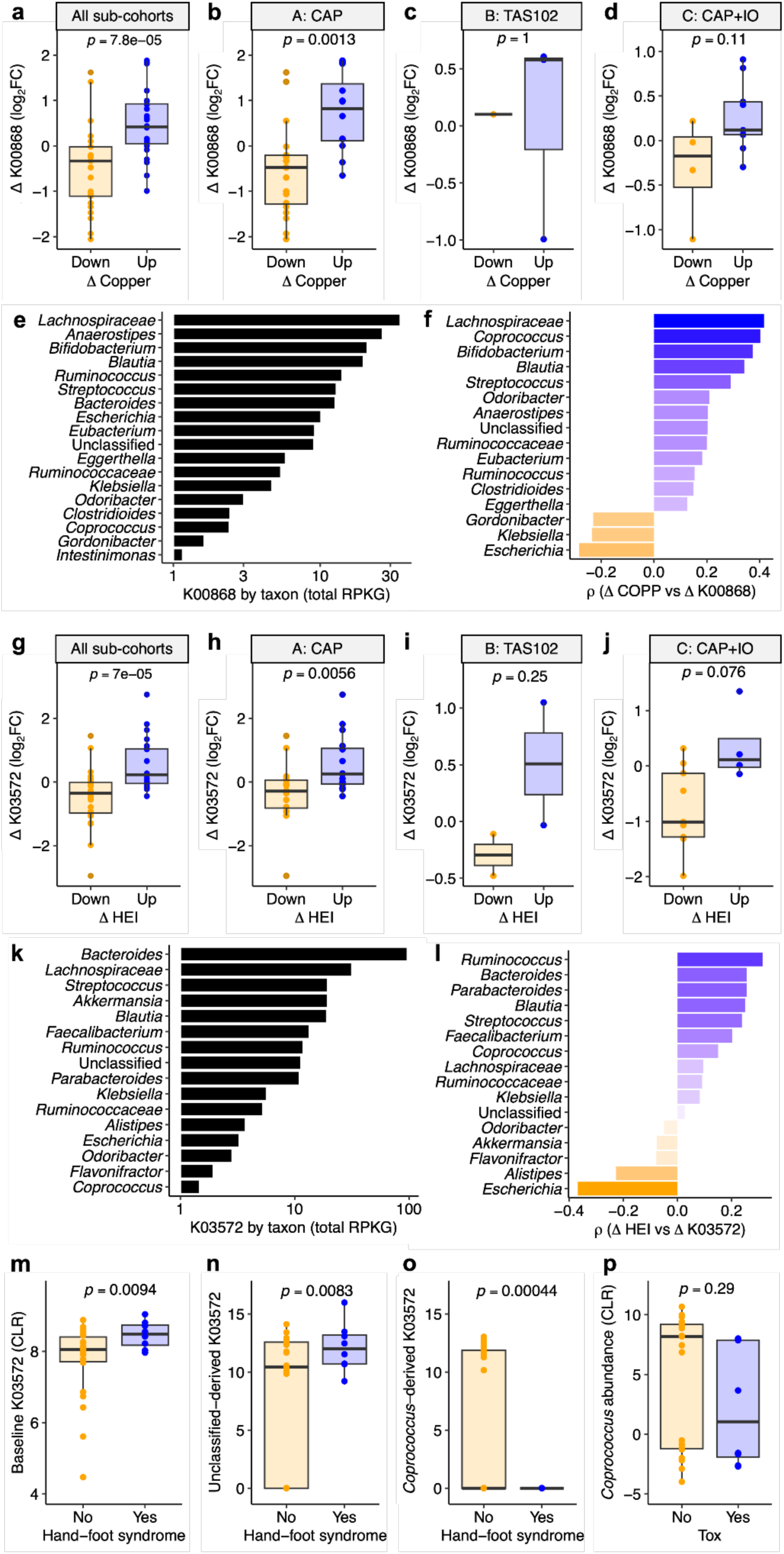
Dietary intake shifts correlate with micronutrient- and chemotherapy-related gut microbial genes. (**a-d**) Change in K00868 (*pdxK*) gene abundance vs change in copper intake. Sub-panels represent data from all subjects (**a**) or stratified by sub-cohort (**b-d**). (**e**) Genus-level contribution to *pdxK* abundance. (**f**) Genus-level *pdxK*-copper correlations. (**g-j**) Change in K03572 (*mutL*) gene abundance vs change in HEI. Sub-panels represent data from all subjects (**g**) or stratified by sub-cohort (**h-j**). (**k**) Genus-level contribution to *mutL* abundance. (**l**) Genus-level *mutL*-HEI correlations. (**m-p**) Development of hand-foot syndrome vs total baseline abundance of K03572 (**m**), K03572 of Unclassified genus source (**n**), *Coprococcus*-derived K03572 (**o**), and *Coprococcus* genus abundance (**p**). *p*-values: (**a-d, g-j**) Mann-Whitney U test, where each point represents the difference between 2 consecutive timepoints within a single patient; (**f**,**l**) Spearman correlation between change in dietary intake (log₂ for copper, raw for HEI) vs log₂ fold-change gene abundance between each pair of consecutive within-patient timepoints; (**m-p**) Welch’s *t*-test.

**Extended Data Figure 8:**
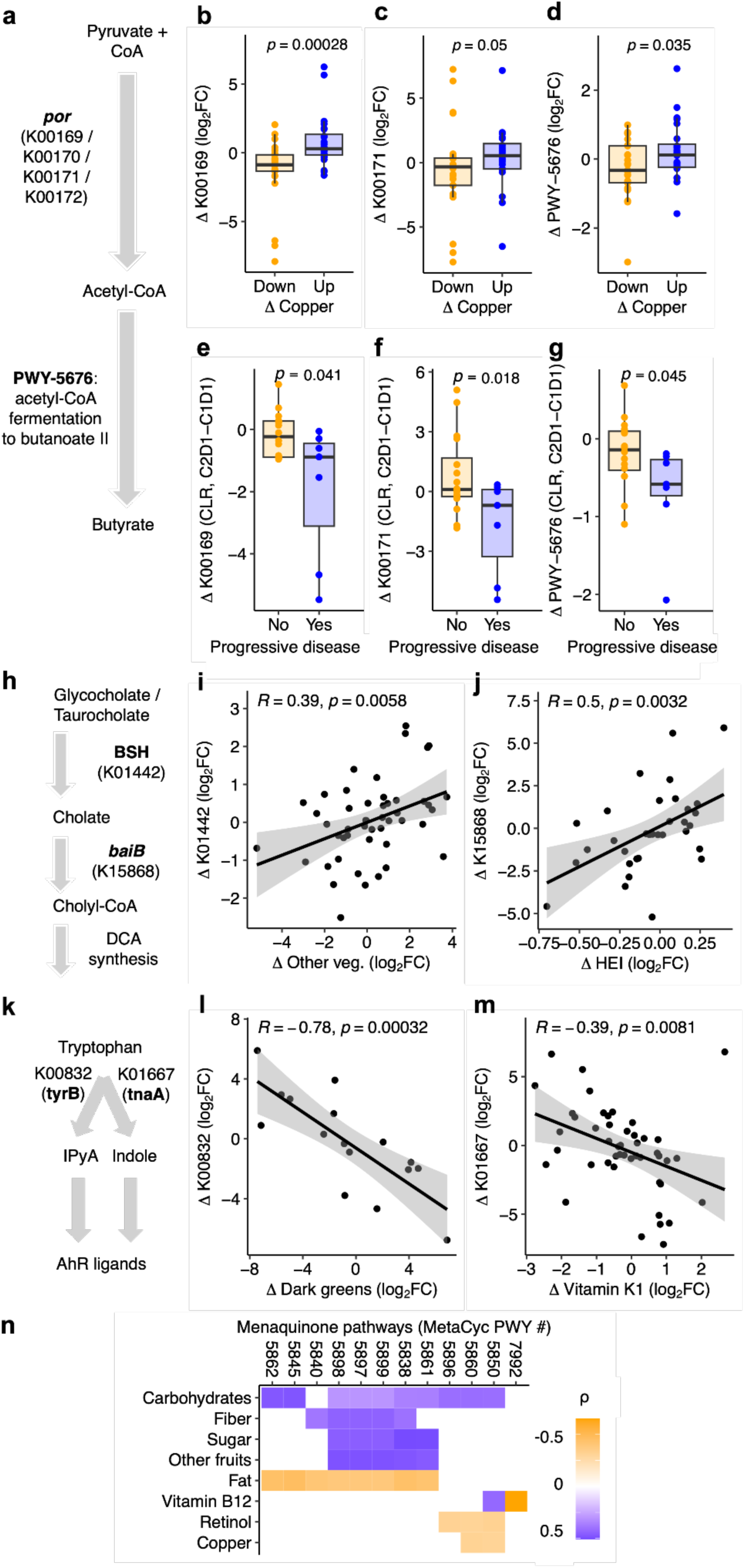
Dietary intake shifts correlate with CRC-related microbial gene functions. (**a**) Microbial production of butyrate from pyruvate, adapted^26,27^. (**b,c**) Change in butyrate pathway pyruvate:ferredoxin oxidoreductase K00169 (*porA*, **b**) and K00171 (*porD*, **c**) gene abundance vs change in copper intake. (**d**) Change in PWY-5676: acetyl-CoA fermentation to butanoate vs change in copper intake. Despite the positive PWY-5676-copper association, no individual genes in PWY-5676 were significantly associated with copper dynamics (FDR>0.2; Fig. 4a**,b**, **Table S10**). (**e-g**) Change in butyrate gene abundance during treatment vs disease progression: *porA* (**e**); *porD* (**f**); PWY-5676 (**g**). (**h**) Microbial bile acid synthesis pathway, adapted^28^. (**i-j**) Change in BSH (K01442, **i**) and *baiB* (K15868, **j**) vs change in other vegetable intake (**i**) and HEI (**j**) between each pair of consecutive within-patient timepoints, respectively. No other dietary components were positively associated with K01442 or K15868 (FDR>0.2; Fig. 4a). (**k**) Schematic of initial step of microbial tryptophan metabolism to AhR ligands^29^. IPyA: indole-3-pyruvic acid. (**l-m**) Change in *tyrB*/*araT* (K00832, **l**) and *tnaA* (K01667, **m**) vs change in other dark greens (**l**) and vitamin K_1_ (**m**) intake between each pair of consecutive within-patient timepoints, respectively. Starchy vegetable dynamics also negatively correlated with K00832 (R = -0.66, *p* = 0.0023); no other dietary features were linked to K00832 or K01667 (FDR>0.2; Fig. 4a). (**n**) Because vitamin K_2_ biosynthesis involves many genes and is well captured by HUMAnN pathway definitions, we focused on pathway-level rather than KO-level associations. Heatmap of change in menaquinol pathways vs change in dietary intake. Color depicts ρ magnitude and sign; all colored tiles have FDR<0.2. *p*-values: (**b-g**) one-tailed Mann-Whitney U test; (**i,j,l,m**) linear regression of log₂ fold-change of diet intake vs log₂ fold-change microbiome feature abundance, with solid black line indicating the line of best fit and grey bands the 95% confidence interval; (**n**) Spearman correlation of change in rank-transformed dietary intake vs change in rank-transformed central log ratio [CLR]-normalized HuMANN pathway abundance; all colored tiles have FDR<0.2, where Benjamini-Hochberg false discovery rate (FDR) correction was performed across all tested genes for each dietary variable.

**Extended Data Figure 9:**
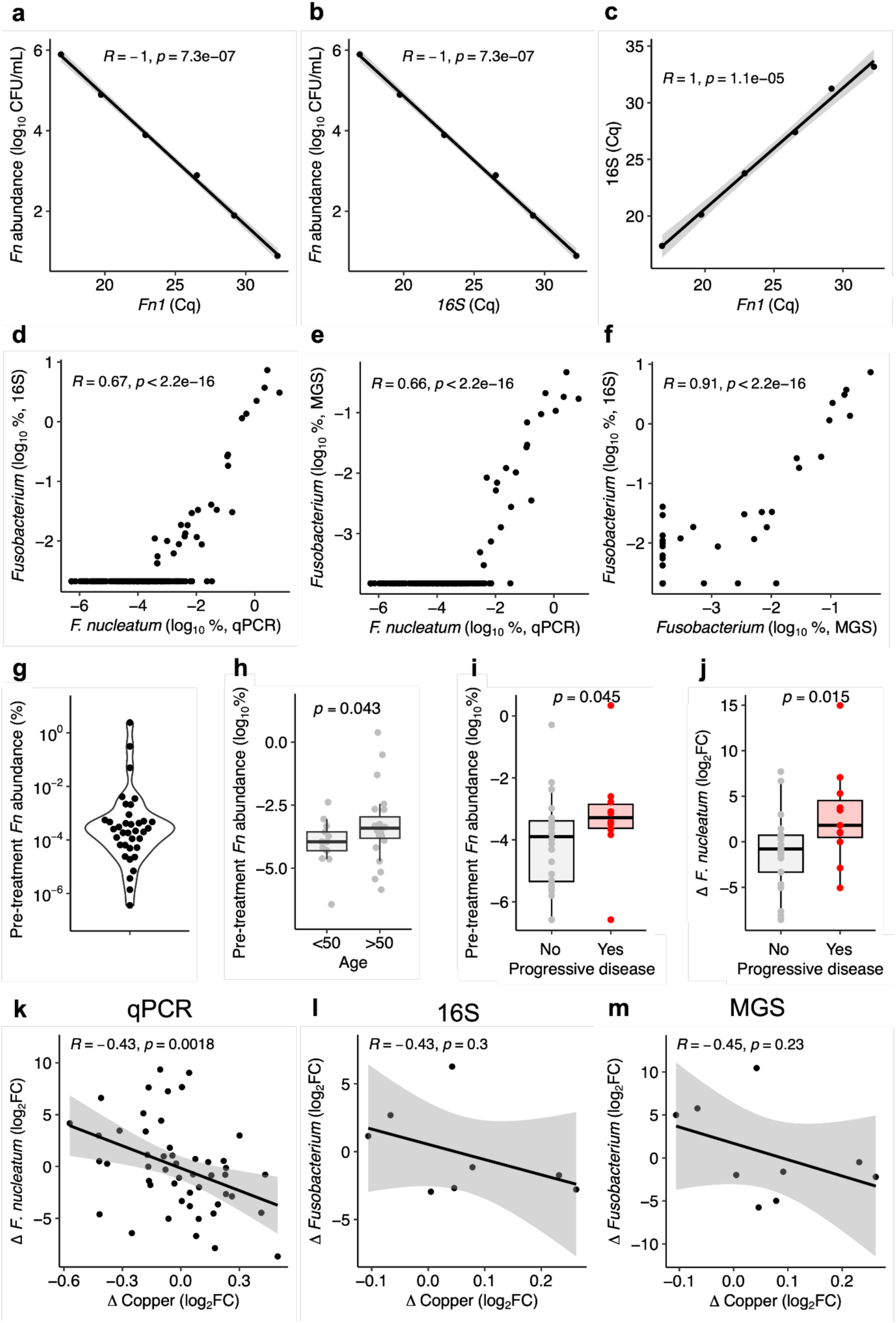
Decreased copper intake correlates with decreased *Fusobacterium*. (**a,b**) Dilutions of gDNA extracted from *Fn* DSM15643 pure culture vs Cq of *Fn*-specific 16S region *Fn1* (**a**) or general 16S (**b**). (**c**) *Fn1* Cq vs general 16S Cq for pure culture of *Fn* DSM15643. (**d-f**) Pairwise correlations of *Fusobacterium* abundance estimated from qPCR (*F. nucleatum*), 16S (*Fusobacterium* genus), and metagenomic sequencing (MGS; *Fusobacterium* genus). (**g**) Per-patient pre-treatment (C1D1) *F. nucleatum* abundance. (**h**) Pre-treatment *F. nucleatum* abundance vs young-onset CRC status (yoCRC, age < 50). (**i,j**) Pre-treatment *F. nucleatum* (**i**) and change in *F. nucleatum* abundance (C2-C1, **j**) vs progressive disease status. (**k-m**) Change in copper abundance vs change in *Fusobacterium* abundance between each pair of consecutive within-patient timepoints where at least 1 timepoint has *Fusobacterium* detected, using qPCR (**k**), 16S (**l**), and MGS (**m**) to estimate *Fusobacterium* abundance. *p*-values: (**a-f, k-m**) Pearson correlation, where solid black indicates the line of best fit, and grey bands indicate the 95% confidence interval; (**h-j**) one-tailed Mann-Whitney U test.

**Extended Data Figure 10:**
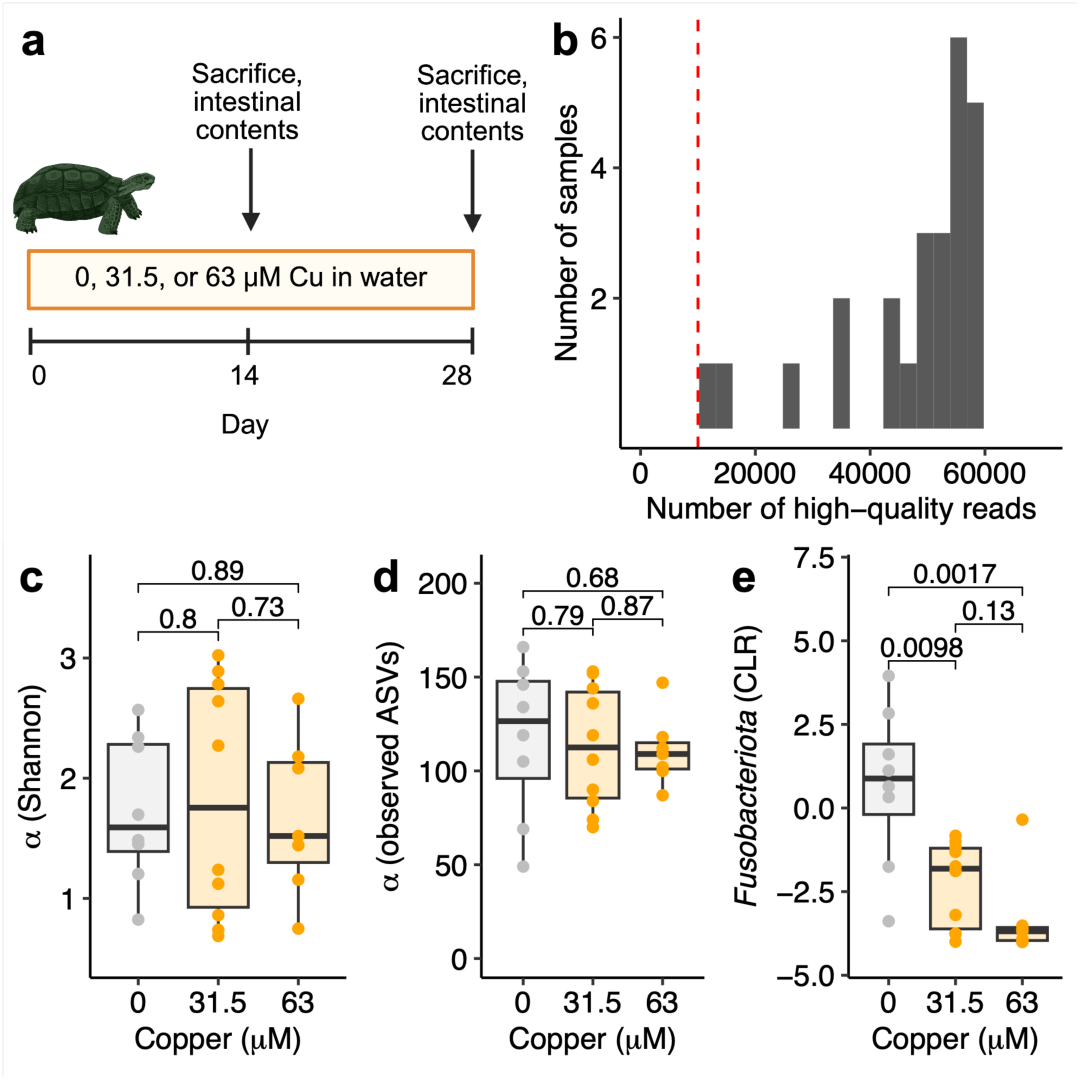
Copper suppresses *Fusobacteriota* in vivo. Re-analysis of raw sequencing data from Ali *et al*^34^. (**a**) Study design. *Mauremys sinensis* turtles were treated with 0, 31.5, or 63 μM CuSO_4_ for 14 or 28 days, followed by sacrifice and collection of pan-intestinal contents. Created with BioRender. (**b**) High-quality read histogram. The red dotted line represents a high-quality read threshold of 10,000, which all samples exceeded. (**c-d**) Alpha diversity as measured by Shannon index (**c**) and number of observed ASVs (**d**) vs copper. (**e**) *Fusobacteriota* centered log-ratio (CLR) abundance vs copper. *p*-values: Welch’s *t*-tests (*n*=7-9/group).

## SUPPLEMENTAL FIGURES AND FIGURE LEGENDS

**Figure S1:**
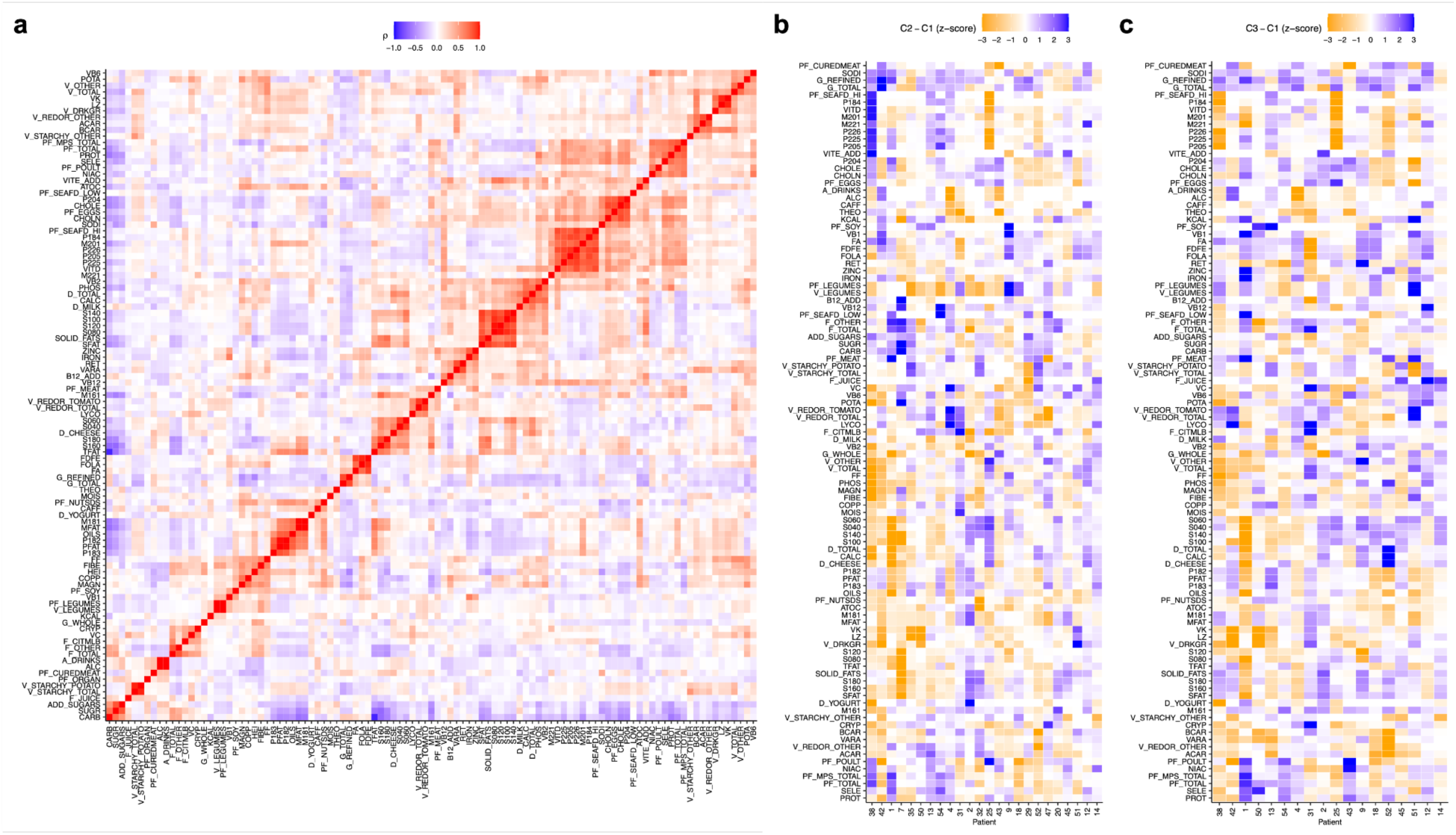
Patterns of nutrient co-variation and temporal dynamics. (**a**) Pearson correlation between all pairs of dietary variables. Highly correlated variables (ρ>0.85) were dereplicated. (**b,c**) Per-patient change in nutrient intake (z-score difference) between cycle 1 (C1) and cycle 2 (C2) (**b**), and between C1 and cycle 3 (C3) (**c**).

**Figure S2:**
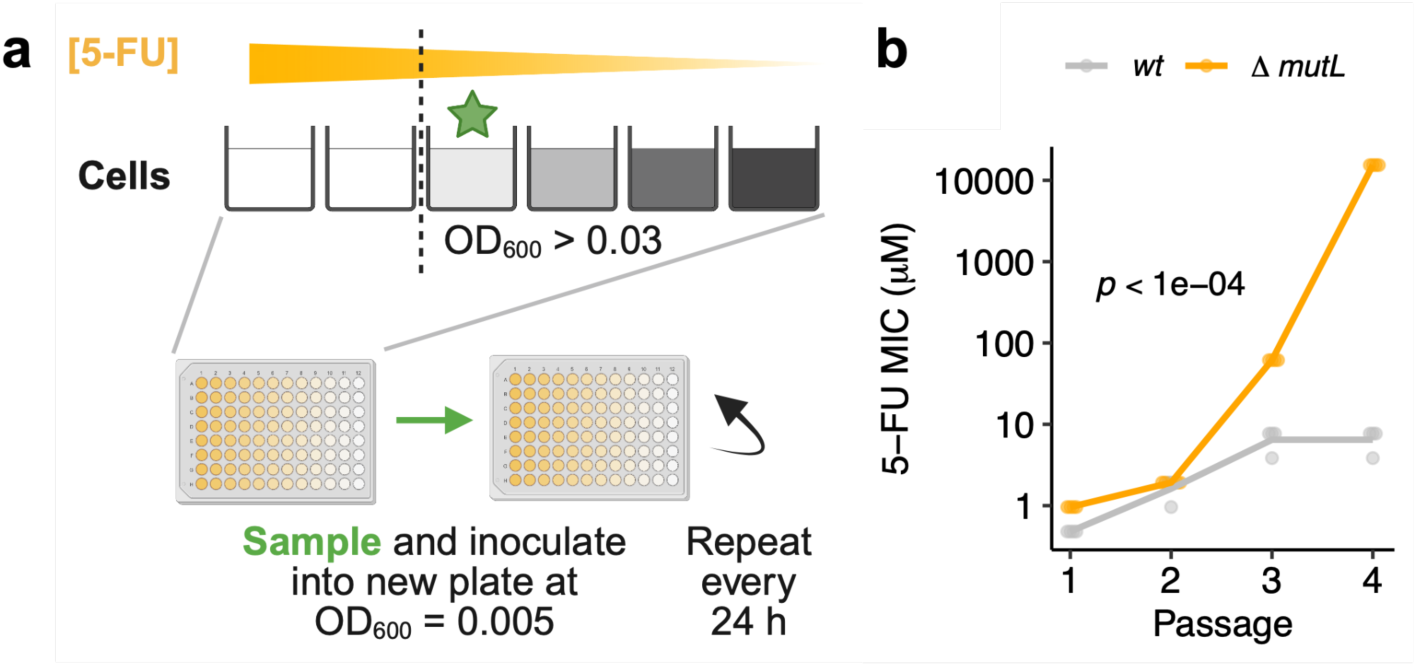
Experimental evolution under 5-FU selection pressure. **(a)** Experimental design (made with BioRender, inspired by Ref^85^). A 96-well plate was prepared with 2-fold serial dilutions of 5-FU in M9-Glc. After 24 hours, cells from the well with the highest drug concentration with sufficient cell growth (OD_600_ > 0.03) were transferred to fresh media with a 5-FU gradient. Serial transfers were performed over 4 days. **(b)** 5-FU MIC of E. coli BW25113 *wt* and *ΔmutL* serially passaged for 24 hours in M9-Glc ± varying concentrations of 5-FU as part of the experimental evolution experiment from **(a)**. *mutL* deletion accelerates evolution of 5-FU resistance. *p*-value: strain*passage interaction term from two-way ANOVA.

**Figure S3:**
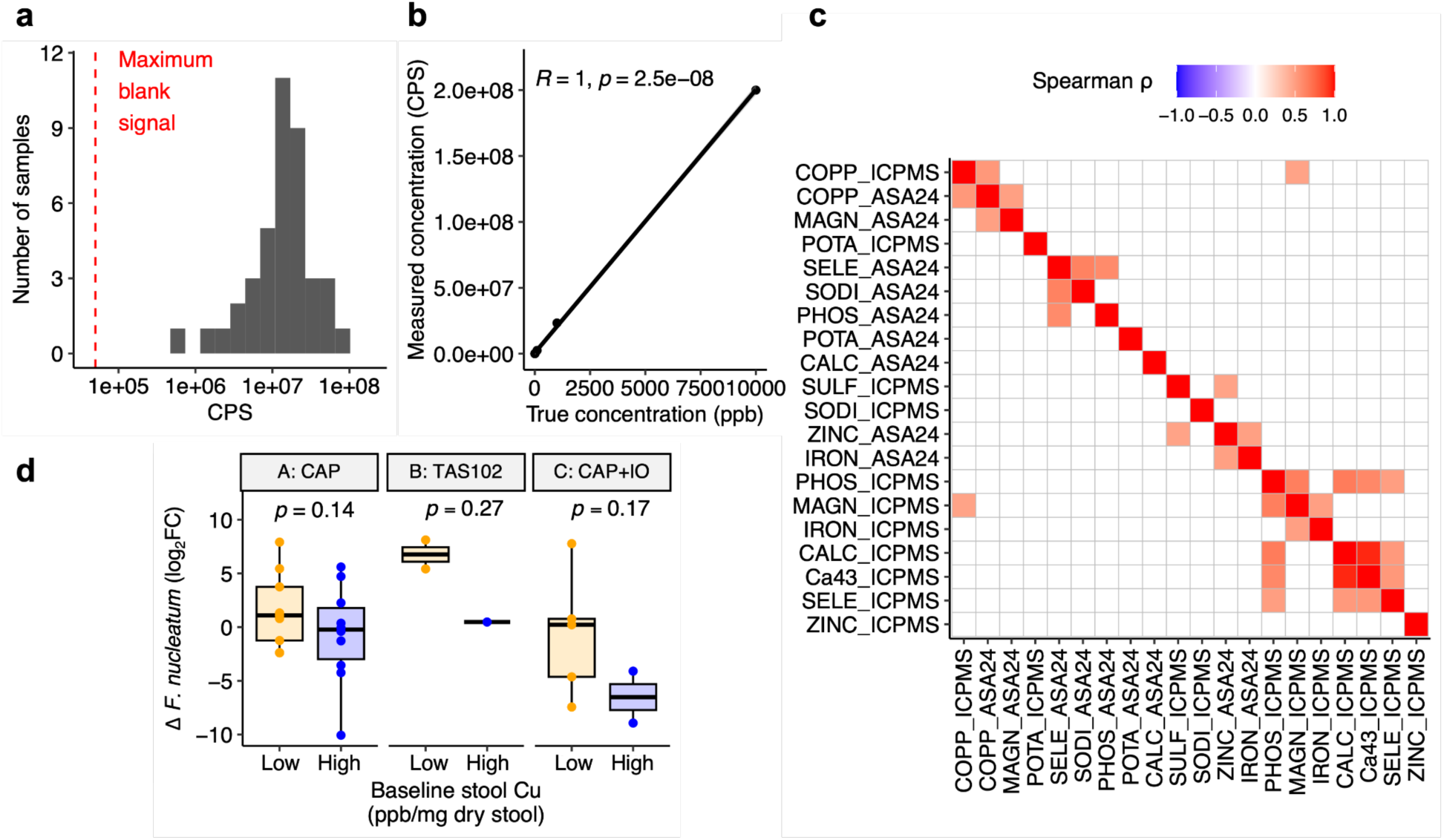
Measurement of stool ion concentrations. (**a**) Distribution of raw copper signal for each stool sample (counts per second, CPS). (**b**) Copper standard curve. *p*-value: Pearson correlation. (**c**) Autocorrelation of dietary metal intake (ASA24) and stool metal concentration (ICPMS). *p*-value: Spearman correlation. Only tiles with *p* < 0.05 are colored. (**d**) Change in *F. nucleatum* abundance during cycle 1 vs baseline stool copper levels, stratified by sub-cohort. Copper is called Low/High relative to the median of the entire dataset. *p*-value: one-sided Welch’s *t*-test.

**Fig. S4:**
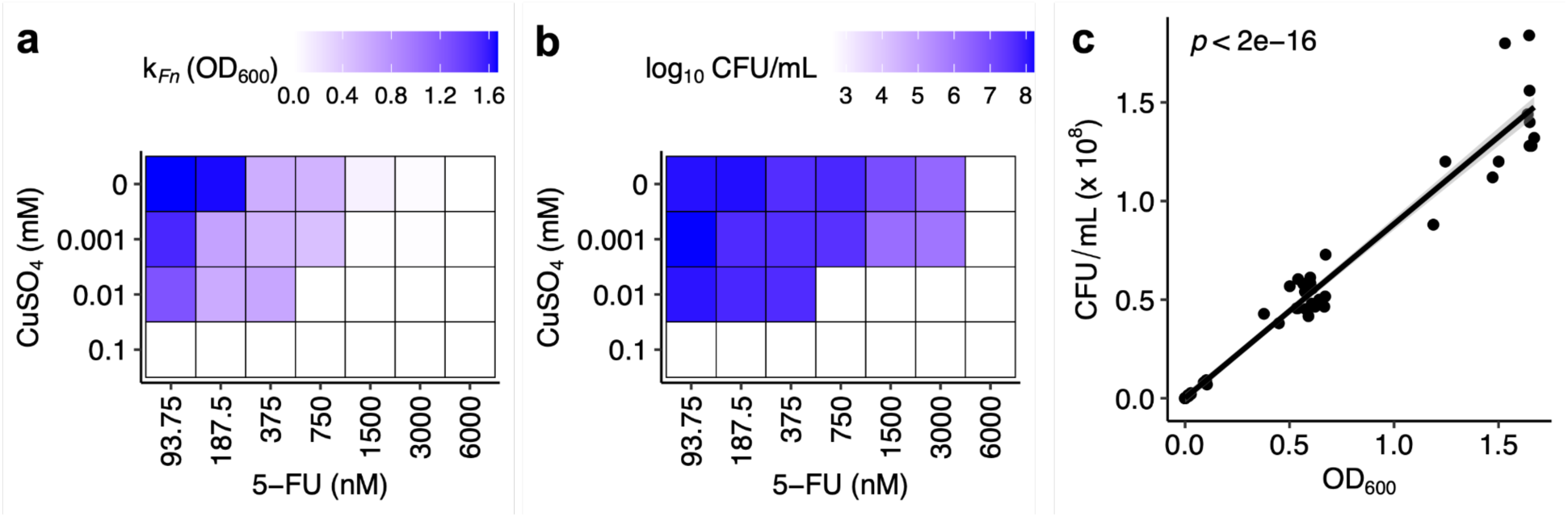
Copper suppresses *Fusobacterium nucleatum* in vitro. *F. nucleatum (Fn)* was grown for 24 hours in BHI^CHV^ ± CuSO_4_ (0, 0.001, 0.01, 0.1 mM) and 5-FU (6000, 3000, 1500, 750, 375, 187.5, 93.75 nM) (*n*=2-4 replicates pooled from 2 independent experiments), with growth assessed by optical density (OD_600_; carrying capacity k*_Fn_*) (**a**) and colony-forming units (CFU) (**b**). (**c**) Correlation of endpoint optical density (OD_600_) and colony counts (CFU/mL x 10^8^) for each replicate from (**a,b**). *p*-value: Spearman correlation. The solid black is the line of best fit, and grey bands indicate the 95% confidence interval. For (**b,c**), values below the limit of detection of 400 CFU/mL were set to 400.

